# Deep learning-based codon optimization with large-scale synonymous variant datasets enables generalized tunable protein expression

**DOI:** 10.1101/2023.02.11.528149

**Authors:** David A. Constant, Jahir M. Gutierrez, Anand V. Sastry, Rebecca Viazzo, Nicholas R. Smith, Jubair Hossain, David A. Spencer, Hayley Carter, Abigail B. Ventura, Michael T. M. Louie, Christa Kohnert, Rebecca Consbruck, Joshua Bennett, Kenneth A. Crawford, John M. Sutton, Anneliese Morrison, Andrea K. Steiger, Kerianne A. Jackson, Jennifer T. Stanton, Shaheed Abdulhaqq, Gregory Hannum, Joshua Meier, Matthew Weinstock, Miles Gander

**Affiliations:** Absci Corporation, Vancouver (WA) and New York (NY), USA

**Author notes:** Equal contribution (David A. Constant, Jahir M. Gutierrez, Anand V. Sastry, Rebecca Viazzo, Nicholas R. Smith).

## Abstract

Increasing recombinant protein expression is of broad interest in industrial biotechnology, synthetic biology, and basic research. Codon optimization is an important step in heterologous gene expression that can have dramatic effects on protein expression level. Several codon optimization strategies have been developed to enhance expression, but these are largely based on bulk usage of highly frequent codons in the host genome, and can produce unreliable results. Here, we develop deep contextual language models that learn the codon usage rules from natural protein coding sequences across members of the *Enterobacterales* order. We then fine-tune these models with over 150,000 functional expression measurements of synonymous coding sequences from three proteins to predict expression in *E. coli*. We find that our models recapitulate natural context-specific patterns of codon usage and can accurately predict expression levels across synonymous sequences. Finally, we show that expression predictions can generalize across proteins unseen during training, allowing for *in silico* design of gene sequences for optimal expression. Our approach provides a novel and reliable method for tuning gene expression with many potential applications in biotechnology and biomanufacturing.

## Introduction

Industrial production of recombinant proteins is a major component of biomanufacturing with a wide range of applications [1]. As of 2018, there were 316 licensed protein-based biopharmaceutical products with sales totaling at least $651 billion since 2014 [2]. Microbial systems such as *E. coli* [3] have long been a workhorse of recombinant protein production, and are important biopharmaceutical manufacturing platforms with several advantages over conventional mammalian cells, such as scalability and affordability [4, 5]. As of 2022, there were at least 29 biopharmaceuticals produced in *E. coli* [6]. Thus, increasing production efficiency by boosting protein expression in microbial hosts can have a significant impact on the affordability and availability of pharmaceuticals and other biomanufactured products [7].

The expression level of recombinant proteins depends on multiple factors, including associated regulatory elements flanking the gene coding sequence (CDS) [8, 9], culture conditions used for growing the production host cells [10], metabolic state of such cells [11], or co-expression of chaperones [12]. Particularly, codon usage in the CDS is an important factor that has been exploited to increase recombinant protein expression in biotechnology and synthetic biology [13–16]. Codon usage patterns have been shown to affect expression via changes in translation rates, mRNA stability [17], protein folding, and solubility [18, 19]. Today, several commercial tools and algorithms exist to optimize codon usage in a CDS. These typically rely on sampling codons from the distribution of frequently observed codons in the host genome [19–23], maximizing the codon adaptation index (CAI) or codon-pair biases [24, 25]. However, these tools fail to account for long-range dependencies and complex codon usage patterns that arise in natural DNA sequences and do not reliably produce high expression CDSs [26–28]. For example, existing codon optimization tools may yield suboptimal DNA sequences that transcribe well in the host, but impede proper folding of the recombinant protein during translation [29]. Alternatively, existing tools may yield sequences that enable proper folding but limit the stability and expression level of the transcribed mRNA [30–34], resulting in diminished yields of functional, soluble protein. For these reasons, developing novel codon optimization strategies capable of capturing these complex and long-range dependencies in DNA sequences is of high interest. Further, establishing language models that can predict DNA sequences and associated expression level for a given host will be a major step towards understanding the underlying principles governing gene expression.

Natural language models based on Deep Learning (DL) have emerged as powerful tools for interrogating complex context-dependent patterns in biological sequences across application domains [35–38]. Although there are recent examples of DL-enabled codon optimization [39–41], these studies do not incorporate expression level information. Here we show that language models are able to learn long-range patterns of codon usage and generate sequences mimicking natural codon usage profiles when trained on genome-scale CDS data. We find that optimizing gene sequences for natural codon usage patterns alone does not guarantee high protein expression level; rather, additional optimization based on functional expression level data is necessary to reliably identify gene sequences with high expression.

We extend the use of language models for codon optimization by predicting protein expression levels via training on a large collection of paired sequence to expression data for three distinct recombinant proteins. These functional protein expression datasets were generated via multiple assays including our previously described Activity-specific Cell Enrichment (ACE) assay [37, 42], a Sort-seq method [16] and antibiotic selection. Taken together, the full dataset accounts for 154,166 total functional expression measurements of synonymous full gene sequences and, to our knowledge, represents the largest dataset of its kind. With this dataset, our trained models predict expression level of unseen sequence variants with high accuracy. Finally, we predict and experimentally validate high-expressing sequences for two proteins outside the model training set, demonstrating the generalized ability for our model to design CDSs for optimal functional expression of proteins.

## Results

### Generative deep language models optimize CDSs for natural sequence attributes but do not reliably create high expressing variants

One common approach for codon optimization is a text translation task, where a sentence written in the language of protein amino acids is translated to DNA nucleotides, conditioned on the identity of the host organism to maximize the natural profile of the CDS [39–41]. To this end, we repurposed the T5 language model architecture [43] by training on a dataset, named Protein2DNA, consisting of all protein coding and protein-CDS pairs from high-quality genomes of the *Enterobacterales* order (taxonomic identifier 91347) (Materials and methods). We trained our model, referred to as CO-T5, on the single task of protein-to-DNA translation, conditioned on the taxonomy of the host genome. Training in this novel way allows the model to learn rich, shared gene sequence patterns across organisms. By providing the identity of the host organism, CO-T5 learns to output CDSs specific to each taxonomy. Our model represents the first deep generative language model for full-length CDSs.

To test CO-T5’s ability to learn natural sequence patterns we generated DNA sequences from a holdout set of proteins from various members of the *Enterobacterales* order, and compared the generated sequences to their endogenous versions. As shown in figure 1A, generated sequences and their natural counterparts have similar Codon Adaptation Index (CAI) [44], a measure of host codon frequency in a CDS. To determine if model-generated sequences have natural long-range codon usage patterns we also investigated the %MinMax profiles [26] across generated sequences, compared to their natural counterparts. %MinMax measures the prevalence of rare codons in a sliding window, where lower values indicate clusters of rare codons. For comparison, we generated synonymous sequences for each natural amino acid sequence by randomly sampling codons at each position. The %MinMax profiles of CO-T5-generated sequences are remarkably similar to their natural counterparts (fig. 1B,C) compared to random degenerate sequences, demonstrating the model’s ability to recapitulate DNA sequences with natural attributes. We also computed sequence similarities (fig. 1D) and found that CO-T5-generated sequences, from holdout examples, have an average sequence similarity of 85% to their natural counterparts in contrast to an average of 75% for random synonymous sequences.

**Fig 1.**
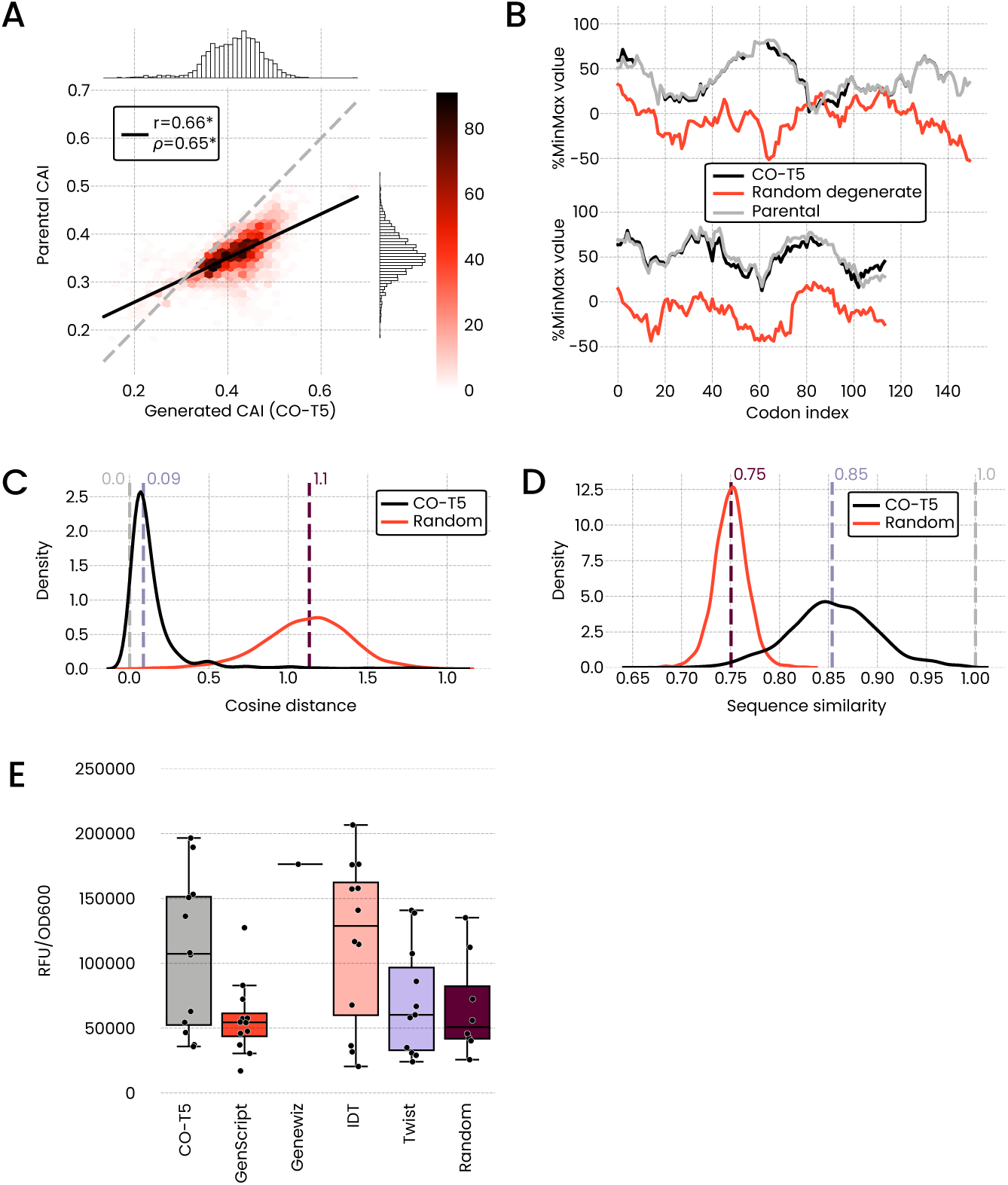
Codon sequences generated by the CO-T5 model recover natural sequence attributes and express at variable levels. **(A)** Heatmap correlation of CAI, a measurement of codon frequency in CDSs, from CO-T5 model-generated and natural sequences in the *Enterobacterales* holdout set. Dashed line represents unity and the solid line is best fit. **(B)** Two representative %MinMax codon usage profiles of CDSs in the holdout test set. The comparison is made across parental natural sequence (grey), CO-T5-model-generated sequence (black), and sequence of randomly sampled synonymous codons (red). **(C)** Kernel density plot of cosine distances between %MinMax profiles in the holdout set for CO-T5-generated (black) and random synonymous (red) sequences versus their natural counterparts. The dashed lines represent the identity with natural counterpart DNA sequence (grey), the average distance for model-generated sequences (lavender), and the average distance for random synonymous sequences (maroon). **(D)** Kernel density plot of DNA sequence similarity in the same holdout set for model-generated and random synonymous sequences versus their natural counterparts. The color legend is the same as in (C). **(E)** Normalized fluorescence values for CO-T5-generated GFP variants, sequences optimized with commercial tools, and random synonymous sequences. Note the Genewiz algorithm is deterministic and only returns a single CDS per input protein.

Previously, we introduced the concept of an antibody Naturalness score [37] to denote how natural a given antibody sequence is according to a pre-trained language model. Here, we introduce the concept of a codon Naturalness score, which is a model-generated score of how natural a codon sequence is in the context of the host genome. Formally, the codon Naturalness score of a CDS is the inverse of the pseudo-perplexity value computed by the CO-T5 language model. We tested whether the natural-like generated DNA sequences from our model were associated with high expression, and compared them to other commonly used optimization algorithms [20–23]. We expressed sequences generated by our CO-T5 model with high Naturalness scores, sequences designed with commercially available codon optimizers [20–23], and random synonymous sequences (fig. 1E). While the CO-T5-generated sequences were among the highest expressors, no statistically significant difference was found by one-way ANOVA between CO-T5 and commercial algorithms, except GenScript (p*<*0.05, Materials and methods). Showing that highly natural CDSs do not necessarily express highly, likely due to natural expression variation across genes [45]. Based on these findings, we next attempted to use supervised learning and fine-tuning to create language models that accurately associate specific codon sequences with expression levels, with the aim of identifying high producing CDSs.

### Masked-language models learn to map CDS to expression levels across a synonymous mutant dataset

Given the varied expression levels of CO-T5-model-generated, high Naturalness score sequences, we devised a supervised learning approach to map specific codon sequences to expression values. To build models that can learn this mapping, we first pre-trained a masked language model called CO-BERTa using the RoBERTa [46] architecture on the same *Enterobacterales* dataset used for CO-T5. We then collected three large-scale datasets of functional expression values for synonymous codon sequences using three different recombinant proteins. These data were used to fine-tune CO-BERTa for the task of predicting expression from a given CDS. As many applications necessitate properly folded, soluble proteins [47], we focused on measuring functional protein levels rather than total protein, although functional assays can often be more difficult to develop.

We used Green Fluorescent Protein (GFP) as our first protein to generate synonymous functional mutant expression data. GFP, a 238 amino acid protein has 2.12 10^110^ (table S1) possible coding variants, many more than is feasible to measure in the laboratory. To focus the search of this massive synonymous mutational space, for effective quantitative screening, we selected three regions, or *tiles*, along the CDS known to affect protein expression (fig. 2A) [15]. For each tile, we constructed a library of degenerate synonymous sequences starting from a parental GFP CDS (Materials and methods, table S2). We then cloned each tile as an independent library of synonymous GFP sequences. This resulted in a highly diverse set of GFP sequence libraries in which only one tile is modified in the CDS at a time (fig. S1A). We then used a Sort-seq method [16] to measure the expression of synonymous GFP sequence mutants in the three libraries (fig. S2-S4). Sort-seq expression measurements were normalized across the libraries (fig. S5-S7) and scaled from 0 to 1, referred to as normalized expression scores. We observed strong correlation (Pearson r = 0.78) between normalized expression scores from Sort-seq and Western Blots of soluble protein for a subset of mutants (fig. S7, S8).

**Fig 2.**
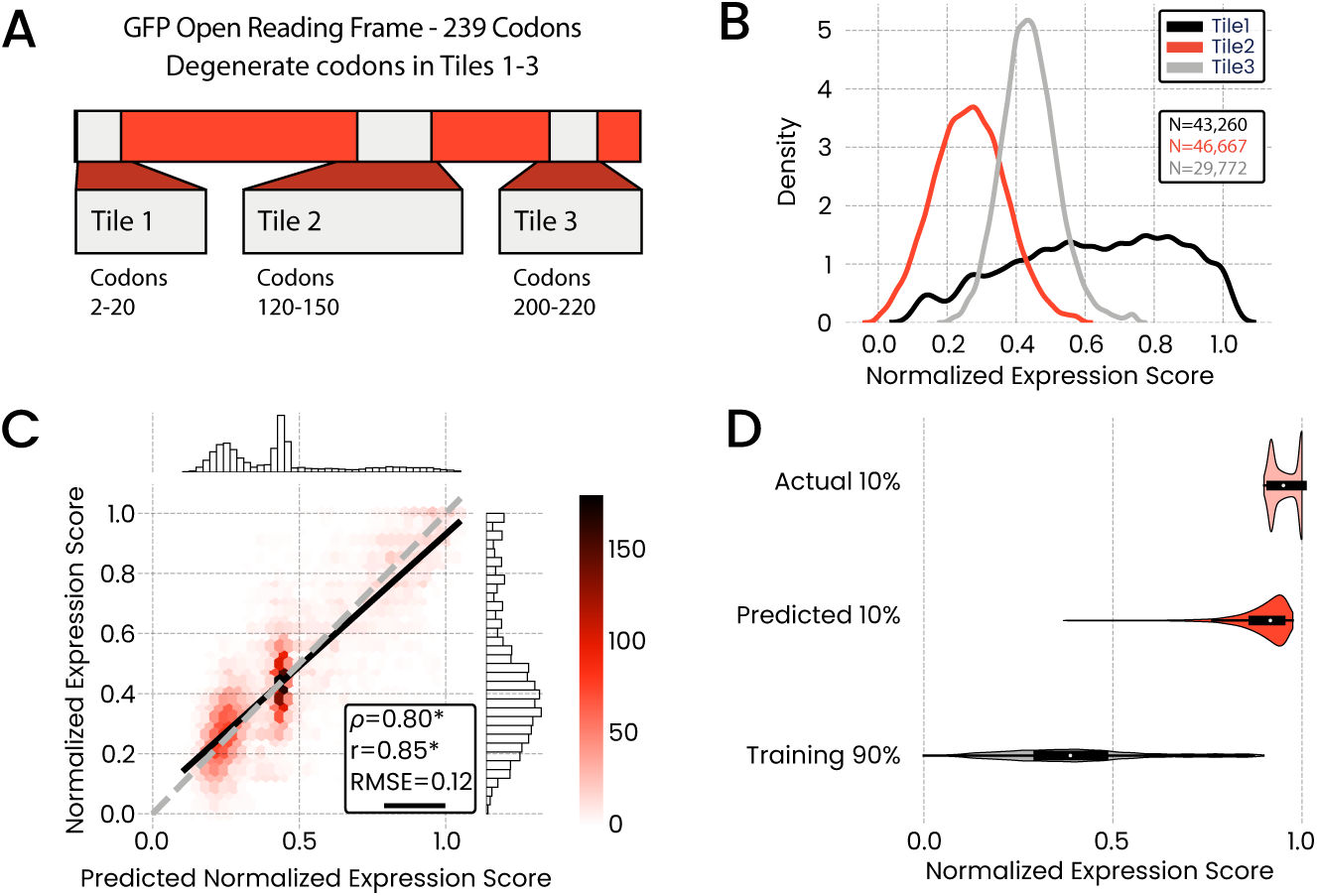
Degenerate synonymous codon GFP libraries. **(A)** Schematic of the three tile synonymous mutant libraries constructed for GFP. **(B)** Density plots of normalized expression scores for all three tile libraries. Distributions show varied expression score profiles based on tile position in CDS. **(C)** Heatmap correlation of GFP model predictions of a test set against measured expression scores. Dashed line represents unity and the solid line is best fit. **(D)** Violin plots depicting model performance of out-of-distribution expression predictions for a holdout set consisting of the top 10% expressing GFP variants. Plot shows bottom 90% variant training set, top 10% variant actual expression measurements, and top 10% expression predictions.

Altogether, the synonymous GFP library included 119,669 measurements of unique CDSs after filtering (Materials and methods, table S3). The distribution of expression levels across these sequences varied according to the position of the tiles, with the first tile, spanning the initial 5’ region of the GFP CDS, having the largest dynamic range and the highest-expressing sequences. This is consistent with previous observations that codon variance in the 5’ region of a given CDS typically has the largest effect on protein expression compared to other regions [15].

The functional GFP data was then used to fine-tune our pre-trained CO-BERTa model. We evaluated its predictive performance, and observed a high correlation between predicted and measured expression levels (fig. 2C; Pearson r = 0.854). Next, we fine-tuned the same pre-trained CO-BERTa model on a capped dataset which held out the highest 10% of measured GFP sequences. We did this to test whether the model could properly generalize to high expression level sequences lying outside of the training distribution. This capped model predicted expression scores similar to the maximal values observed in the training set (fig. 2D), indicating the model’s ability to accurately rank unseen sequences, even if they are outside the range of previously observed variants. Additionally, we used the model to score the GFP variants from figure 1E and observed a correlation between measured fluorescence and predicted expression score (fig. S9; Pearson r = 0.634), further demonstrating the model’s ability to properly score out-of-distribution CDSs. This ranking ability can enable prioritization of high expressors for *in vivo* testing.

### Masked-language models applied to degenerate codon *folA* libraries for expression level prediction

Next we chose the *E. coli* dihydrofolate reductase (DHFR) gene, *folA* for generating synonymous mutant expression measurements. DHFR is a small (~ 18 kDa, 159 amino acids) monomeric enzyme that catalyzes production of tetrahydrofolate from dihydrofolate, and its overexpression confers resistance to drugs inhibiting the folate biosynthesis pathway [48, 49]. The relatively short *folA* CDS enabled use of degenerate oligos to construct a synonymous codon library spanning the entire gene, leading to a highly sequence-diverse library that was bottle-necked into several sub-libraries (fig. S1B).

We used sulfamethoxazole (SMX) and trimethoprim (TMP), synergistic folate biosynthesis inhibitors [50], to select for variants with high expression of DHFR. We observed reduction in cell density at increased levels of antibiotic (fig. 3A) coupled with an increased amount of soluble DHFR production (fig. 3B). We then sequenced post-selection *folA* variants to calculate the frequency of synonymous mutants at increasing concentrations of antibiotic, and computed a weighted average expression score based on sequence prevalence, subjected to score quality filtering (Materials and methods). Again, we normalized and scaled the expression scores from 0 to 1. Replicate selection experiments (N=4) for sub-libraries were highly correlated (fig. S10). During data processing, we observed and filtered out any non-synonymous mutants of DHFR, including those known to increase enzymatic activity (table S3) [51]. We confirmed that our drug resistance functional expression score correlated with soluble expression of DHFR using 24 strains identified in our sequencing results (fig. S11). For these strains, the correlation between dose-response cell density area under the curve (AUC) and protein expression was high for soluble DHFR (Pearson r = 0.884), but low for insoluble DHFR (Pearson r = 0.048), indicating selection for functional, soluble protein. In total, the selection experiments resulted in 17,318 unique sequence variants with associated protein functional expression scores (fig. 3C).

**Fig 3.**
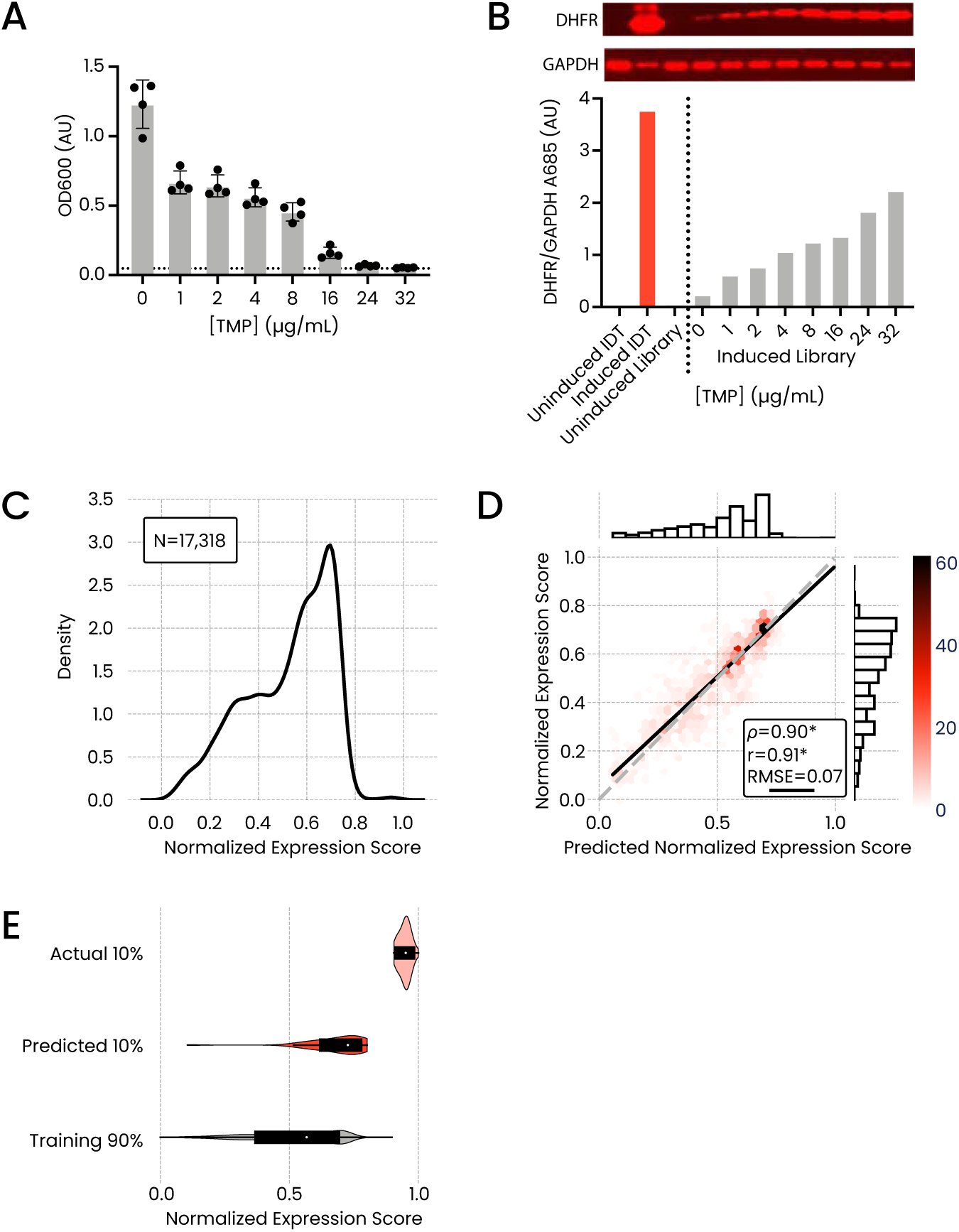
Degenerate synonymous *folA* libraries. **(A)** OD measurements of *folA* degenerate libraries after 24hr of growth at increasing levels of TMP. Plot shows four replicate measurements. Error bars are standard deviation. **(B)** Western Blots of soluble DHFR and GAPDH loading control at increasing levels of TMP for a *folA* degenerate library. Bar chart displays normalized band density values from the blots. **(C)** Normalized expression score distribution from *folA* library selections. **(D)** Heat map correlation of model-predicted expression score values for *folA* variants and expression score measurements for a holdout test set. Dashed line represents unity and the solid line is best fit. **(E)** Violin plots depicting model performance of out-of-distribution expression predictions for a holdout set consisting of the top 10% expressing *folA* variants. Plot shows bottom 90% variant training set, top 10% variant actual expression measurements, and top 10% expression predictions.

Subsequently, we performed model fine-tuning on the *folA* functional dataset. Resulting predictions of the model correlated strongly with measurements (fig. 3D; Pearson r = 0.907). The model also performed well on out-of-distribution predictions of high-expressing sequences (fig. 3E). The top 10% hold out set predictions show a comparably lower normalized expression score than GFP (fig. 1D). Regardless, highly expressing sequences still were predicted with values near the highest within capped 90% training set, demonstrating proper ranking of expression levels for unseen sequences by the model. These results further highlight the ability of model predicted rankings to prioritize testing of high expressing variants in cells.

### Masked-language models applied to a degenerate codon VHH library for expression level prediction

To extend our synonymous codon dataset to include a protein target with relevance to industrial biotherapeutic production, we created a degenerate codon library of an anti-HER2 VHH [52]. VHH-type molecules are heavy-chain-only single-domain antibodies, occurring naturally in Camelid species [53] and can be challenging to functionally express in *E. coli* at high levels due to necessary disulfide bond formation. These molecules are of growing interest as potential biotherapeutics and, thus, increasing their production levels is desirable [54]. We again applied a degenerate tile approach, where the coding parent sequence was altered with either a 5’ degenerate tile, a 3’ degenerate tile, or both degenerate tiles simultaneously. This approach generated a highly diverse library with sequence variation both in focused regions within the tiles in isolation and spanning the whole CDS in dual tile variants (fig. 4A, S1C).

**Fig 4.**
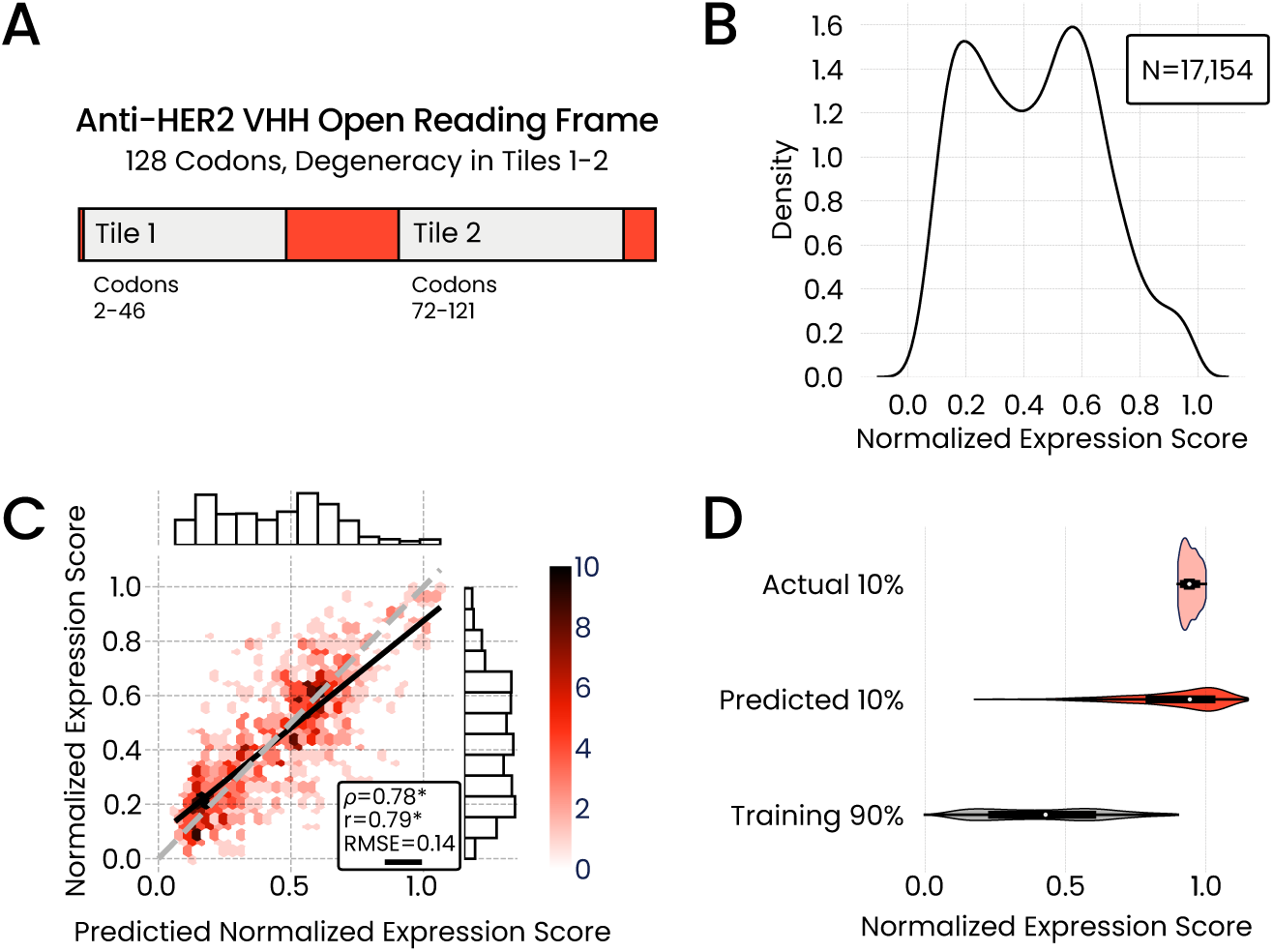
Degenerate synonymous codon VHH libraries. **(A)** Schematic of degenerate tiles for the anti-HER2 VHH parent sequence. **(B)** Normalized expression score distribution of VHH library variants. **(C)** Heatmap correlation of model-predicted VHH expression scores and expression score measurements. Dashed line represents unity and the solid line is best fit. **(D)** Violin plots depicting model performance of out-of-distribution expression predictions for holdout set consisting of top 10% expressing anti-HER2 VHH variants. Plot shows bottom 90% variant training set, top 10% variant actual expression measurements, and top 10% predictions.

We applied a version of the previously described Activity-specific Cell Enrichment (ACE) assay [42] to generate functional protein level measurements for VHH CDS sequence variants (fig. S3B, S12). The ACE assay uses antigen binding fluorescent signal as a proxy for functional protein quantity, coupled with FACS and NGS to generate expression scores for sequence library members. To validate ACE expression scores we performed Western Blots on a subset of VHH sequence variants within the library and found that soluble protein levels were well correlated with ACE expression scores (fig. S13; Pearson r = 0.75). ACE assay screening of the VHH library yielded 17,154 functional expression level measurements of unique CDS variants (fig. 4B).

Next we fine-tuned our models on the VHH dataset and assessed predictive ability similarly to the other protein datasets. We again observed high predictive ability of the model both for in-distribution (fig. 4C) and out-of-distribution sequences (fig. 4D).

### Multi-protein learning

Together, the model performance on all three of the synonymous CDS datatsets demonstrates the ability of language models to learn sequence-to-expression relationships for multiple proteins in isolation. We attempted to improve model performance via multi-task learning across the three proteins. We generated a set of models trained on different combinations of the protein datasets, across either two or all three proteins. We observed an increase in model performance in all cases when training with additional proteins (table 1). Intriguingly, we also observed in some, but not all cases, that models show reasonable performance on proteins outside the training set. The best performance on this unseen protein task was observed with a model trained on *folA* and anti-HER2 VHH, predicting the expression of the GFP dataset. (table S4, S5; Spearman *ρ* = 0.629, Pearson r = 0.558). The improved accuracy in a multi-task training setting and predictive power on some unseen proteins indicates a level of generalizability in the CDS to expression task that could be exploited to design optimized, high expressing DNA sequences.

**Table 1.**
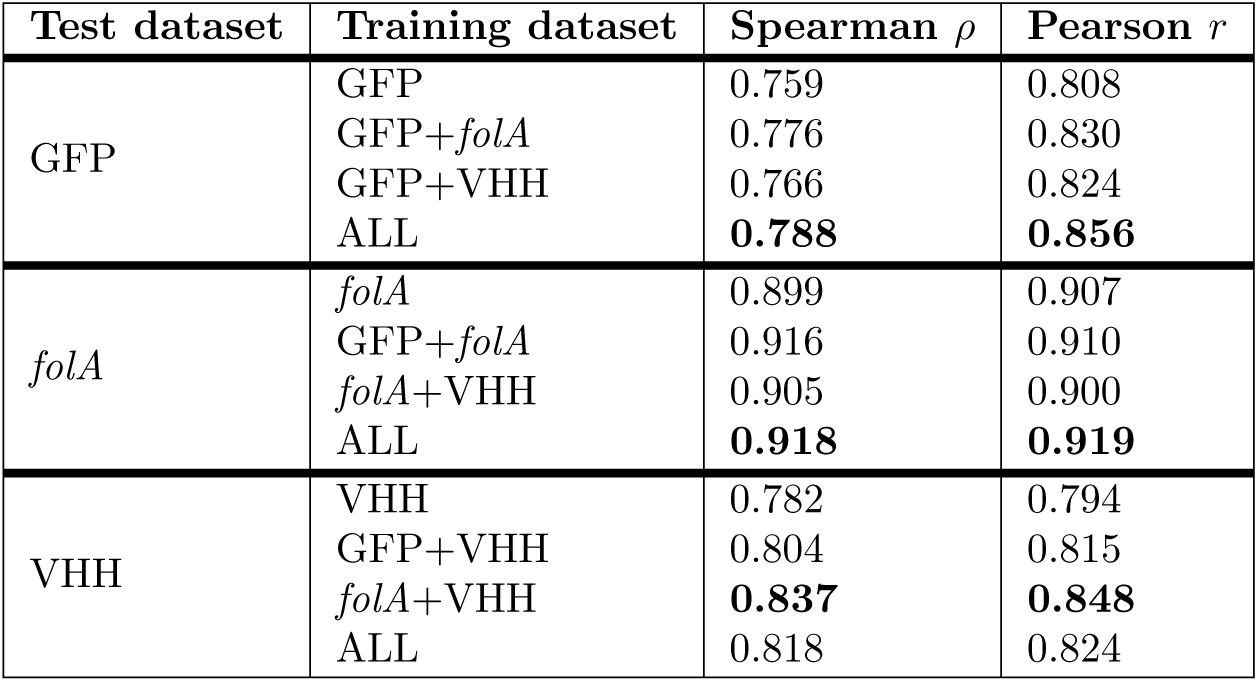
Performance of fine-tuned CO-BERTa models across the GFP (top), folA (middle), and anti-HER2 VHH (bottom) holdout datasets. In all cases we observed increased performance when training on multiple proteins. All p-values across Spearman and Pearson correlations are significant (p<0.01).

| Test dataset | Training dataset | Spearman $\rho$ | Pearson $r$ |
| --- | --- | --- | --- |
| GFP | GFP | 0.759 | 0.808 |
|  | GFP+*folA* | 0.776 | 0.830 |
|  | GFP+VHH | 0.766 | 0.824 |
|  | ALL | **0.788** | **0.856** |
| *folA* | *folA* | 0.899 | 0.907 |
|  | GFP+*folA* | 0.916 | 0.910 |
|  | *folA*+VHH | 0.905 | 0.900 |
|  | ALL | **0.918** | **0.919** |
| VHH | VHH | 0.782 | 0.794 |
|  | GFP+VHH | 0.804 | 0.815 |
|  | *folA*+VHH | **0.837** | **0.848** |
|  | ALL | 0.818 | 0.824 |

### Model baseline comparisons

To further assess our supervised models we performed a number of baseline comparisons (table S4, S5). We created versions of CO-BERTa that were not pre-trained on *Enterobacterales* coding sequences and compared their performance to the pre-trained models from figures 2-4. We find that, in almost all cases, pre-training improves accuracy slightly. Additionally, we compared against traditional Machine Learning (ML) baselines. Specifically, we trained XGBoost models [55] with either (1) the embeddings created by our CO-T5 model or (2) with one-hot encoded representations of the codons (Materials and methods). Interestingly, we find similar performance for boosted tree models on individual proteins to CO-BERTa models. Furthermore, the tree-based models trained on one-hot encoded representations heavily rely on the information provided by the first codons in the sequence to predict expression values (fig. S14-S16), consistent with previous findings [15] and observations in figure 2B. However, training of XGBoost models is constrained to sequences of the same length and, for XGBoost models trained on CO-T5 embeddings, performance does not generalize well across different proteins (table S6). In contrast, our DL models provide the flexibility to train and predict expression level for proteins of different length at higher accuracy. This allows for multi-protein learning which yields a boost in performance (table 1). Additionally, our DL models can generate predictions for unseen proteins, potentially enabling *in silico* design of sequences with specified protein production levels.

### Tuning expression with model-designed DNA variants for unseen proteins

To test the effectiveness of our model as a generalized design tool for modulating protein expression *in vivo*, we created a set of model-designed CDSs for two new proteins outside the training data. We chose mCherry, a monomeric red fluorescent protein, and an anti-SARS-CoV-2 VHH protein [56]. We selected these proteins due to their modest similarity to GFP and the anti-HER2 VHH, respectively, from which we generated our synonymous variant datasets. We hypothesized our models could generalize from the given training set to related proteins. The mCherry protein sequence has 28.7% pairwise identity to GFP, while the anti-SARS-CoV-2 VHH has 73.7% pairwise identity to the anti-HER2 VHH. The new proteins, mCherry and anti-SARS-CoV-2 VHH, differ in amino acid length from their closest counterparts in the training set, GFP and anti-HER2 VHH, respectively (table S2). Despite low sequence identity between GFP and mCherry, the two proteins share a major structural feature, namely a *β*-barrel. Similarly, the two VHH proteins are expected to share high structural concordance. Structural elements can influence codon usage and in turn protein expression and folding [26, 27], potentially enabling our model to generalize to structurally similar proteins outside the training set.

For both mCherry and the anti-SARS-CoV-2 VHH, we designed CDSs with predicted high and low functional expression scores. Sequences were designed through an *in silico* random sampling process via mutating parent CDSs in a tile scheme analogous to the GFP and anti-HER2 VHH libraries (Materials and methods, fig. S17, S18). We iteratively sampled 10^8^ randomized sequences and scored them with the ALL model, trained on all three protein datasets, and either the VHH or full GFP models (table 1). The 10 highest and 10 lowest scored sequences for each protein and model were selected for *in vivo* testing.

We first investigated the functional expression measurements of mCherry variants (fig. 5A). The ALL-model-optimized sequences had the highest mean fluorescence among all conditions and were significantly different by one-way ANOVA from commercial algorithms, excluding the single Genewiz sequence (p*<*0.05). ALL-model-deoptimized sequences showed low expression, near the background of the assay. Interestingly, GFP-model-deoptimized sequences expressed relatively highly, highlighting the benefit of the ALL model’s multi-protein training for generalized expression level tuning of new proteins (fig. S19). To further illustrate the ALL model performance, we calculated the fraction of designed sequences, for each multi-design condition, that fell in the upper quartile of all sequences tested in figure 5A, excluding model-deoptimized designs (fig. 5B). The ALL model outperformed optimization methods capable of generating multiple CDSs by this metric.

**Fig 5.**
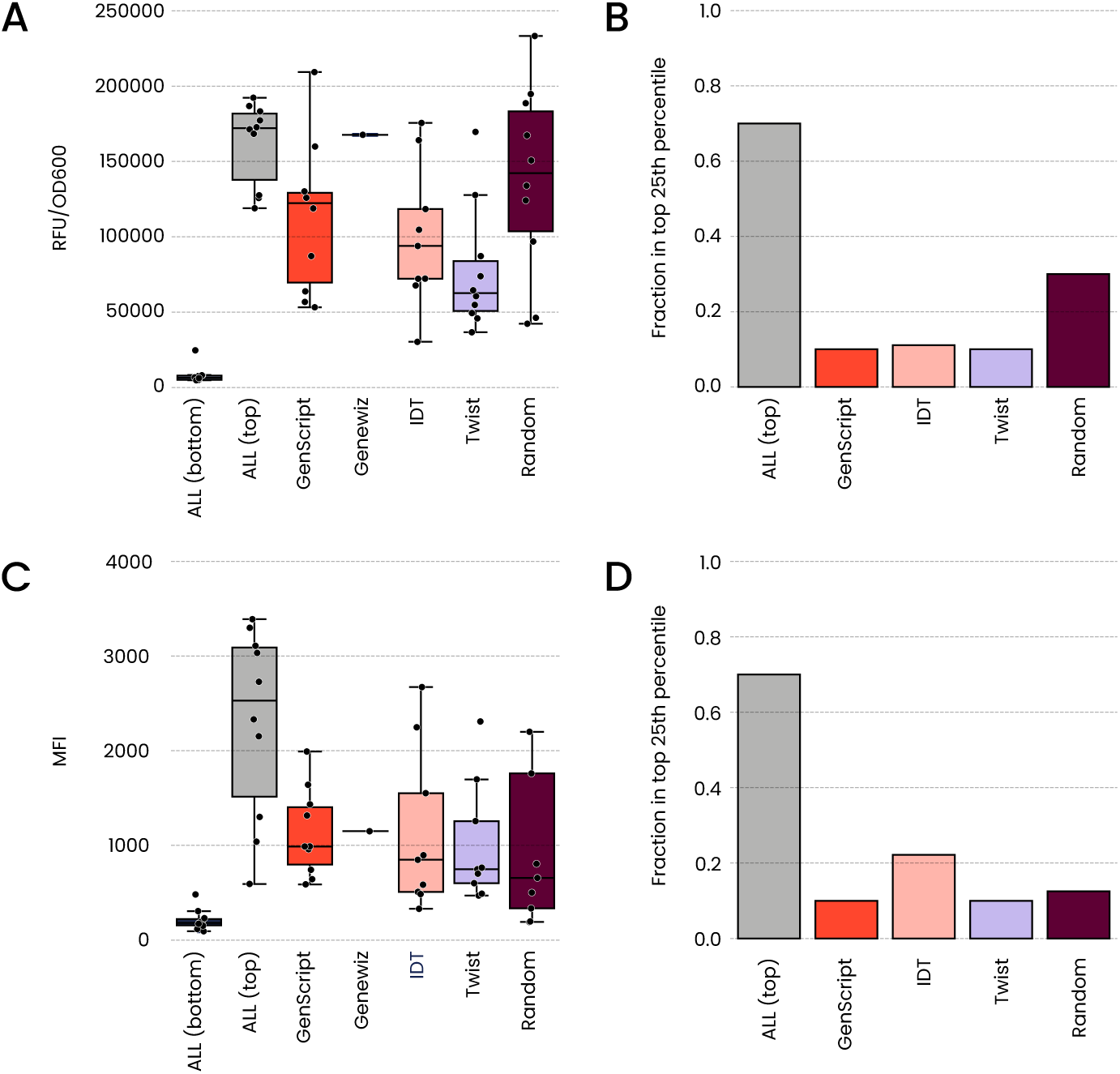
Model-designed protein sequences compared against baselines. **(A)** RFU/OD600 values measured by plate reader for mCherry sequences designed by the ALL model to maximize or minimize expression, compared to various optimization baselines. Values are average of two replicate measurements. **(B)** Barchart shows fraction of designs in upper quartile of all measured mCherry variants for each sequence group, excluding the ALL (bottom) set. Genewiz condition single sequence not included in barchart. **(C)** ACE functional expression measurements, mean fluorescent intensity (MFI), for anti-SARS-CoV-2 VHH sequences designed by the model to maximize or minimize expression, compared to various optimization baselines. Expression values are average of two replicate measurements. **(D)** Barchart shows fraction of designs in the upper quartile of all measured anti-SARS-CoV-2 VHH variants for each sequence group, excluding the ALL (bottom) set. Genewiz condition single sequence not included in barchart.

We performed similar analyses on ACE functional expression level measurements of anti-SARS-CoV-2 VHH variants. We again find the ALL-model-optimized CDSs showed the highest average expression and were significantly different by one-way ANOVA when compared against the commercial algorithms and random sequences, except for Genewiz (p*<*0.05) (fig. 5C). Model-deoptimized sequences once again had low expression levels. We also find the ALL model produced the highest fraction of upper quartile variants from multi-design optimization strategies (fig. 5D). The Genewiz design design did not express in the upper quartile of anti-SARS-CoV-2 VHH variants tested, highlighting the unreliability of this deterministic optimization strategy across multiple proteins. Taken together, these results show the ALL model can be effectively used to design CDSs with specified expression levels for new proteins excluded from the training set.

Additionally, we used the ALL model to score the measured variants in the mCherry and anti-SARS-CoV-2 VHH sets from figure 5. We find a strong correlation between the rankings of expression level and ALL model score (mCherry Spearman *ρ* = 0.72, anti-SARS-CoV-2 VHH Spearman *ρ* = 0.73) (fig. S20). This result shows the ALL model can identify effective candidates for *in vivo* testing, regardless of design method.

## Discussion

Codon optimization for protein expression is a significant challenge mainly due to (1) the large number of possible codon permutations and (2) the complex downstream effects that codons have on processes like transcription [57], translation [58], protein folding [17, 59], and function [60]. For example, mCherry, a 236 amino acid protein, has 1.63 10^107^ possible synonymous DNA sequences. While optimization strategies exist based on codon usage [19], codon pair bias [24], and presence or absence of sequence motifs [13, 15], these approaches lack the ability to capture complex long-range patterns across sequences associated with expression level in this extremely diverse space. Here we demonstrate the ability of deep contextual language models to capture natural codon usage patterns and predict expression level of proteins based on DNA sequence.

We show that while training models on genomic sequence from a given taxon allows for the generation of sequences with natural attributes, the state of the art for DL-enable codon optimization, this alone is not sufficient to consistently generate synonymous DNA sequences with high protein expression levels. To overcome this challenge, we generated the largest ever functional protein expression dataset across three individual proteins and fine-tuned models for protein expression level predictions. Our models can predict CDSs that produce proteins at specified expression levels in the context of a single protein. We also show the model’s ability to accurately rank sequences with protein expression levels higher than observed in the training set, which can save time and resources by prioritizing predicted high-yield variants for *in vivo* testing.

Additionally, we show that training on our functional expression dataset imparts models with the ability to predict expression of DNA variants for proteins outside the training set. Further, this generalizability can be leveraged to design DNA sequence variants with specified expression levels for new proteins. Our model-guided design method outperforms benchmark tools for optimizing and tuning protein expression.

Future work may extend models by increasing the number of diverse protein sequences in training data. Our results suggest that new training examples can increase the accuracy of predictions, which could further improve our generalized model. An additional area of interest would be extending our approach to other organisms beyond *E. coli*. Finally, This work focuses on optimizing protein expression through codon usage, but other studies have used traditional ML or DL techniques to optimize expression via regulatory elements such as promoters [61, 62], ribosome binding sites [63], and terminators [64]. In principle, models accounting for multiple elements involved in expression regulation could be combined to generate a unified model of protein expression.

Taken together, our approach represents a generalized, effective, and efficient method for optimizing the expression of recombinant proteins via codon choice superior to existing methods. Tuning protein expression with our models can have an impact on biomanufacturing and biopharmaceutical availability. We demonstrate the value of applying DL to complex biological sequence space, and provide a framework for increasing protein yield in biological systems.

## Acknowledgments

The authors wish to thank John Hunt, Gŕegory Böel, and Daniel Aalberts for discussion of codon optimization in *E. coli*, Alexandria Kershner for bioinformatics pipeline support, Drew S. Tack for providing the GFP base plasmid, Jia Yeu Liu for productive discussion of *folA* experiments and Sean McClain for his unwavering support. The authors also wish to thank Marcin Klapczynski, Tom Wrona and Stephanie Yasko for providing review, editing and graphic design assistance for the manuscript.

## Materials and methods

### Cloning

For all cloning reactions, backbone fragments were generated by PCR using proof-reading polymerase (Phusion^TM^, ThermoFisher cat #F530 or Q5^®^, NEB cat #M0492). Discrete plasmids were constructed using the HiFi DNA Assembly kit (NEB, cat #E2621) to insert synthetic genes (IDT gBlocks or eBlocks) or isolated as single clones from libraries. All plasmids were verified by sequencing. All thermal cycling conditions are given in supplemental oligo file. Where DNA sequences were optimized using online algorithms, the organism was chosen that was closest to *E. coli* strain B. For IDT, this was *Escherichia coli* B; for Genewiz, GenScript, and Twist, this was *Escherichia coli*. For GFP sequences, All predictions from all algorithms were screened for AsiSI, AscI, BsaI, and BbsI restriction sites. Where optimizers are non-deterministic, the first optimized sequences returned were used with no further filtering.

#### Degenerate GFP libraries

Three regions of the recombinant green fluorescent protein (GFP) nucleotide sequence were chosen for investigation as degenerate libraries (codons 2-20, 120-150, 200-220). Backbone fragments were amplified in two pieces, reactions were treated with DpnI (37 °C, 15 min), and amplicons were gel-purified (Qiaquick Gel Extraction Kit, Qiagen cat #28706) followed by purification (DNA Clean & Concentrator, Zymo Research cat #D4004). Libraries were assembled using the HiFi DNA Assembly kit (NEB, cat #E2621) from degenerate Ultramer^TM^ oligos (IDT). Reactions were assembled with 15 fmol of each backbone fragment and 75 fmol of insert in 20 µL. Reactions were incubated at 50 °C for 60 minutes. For amino acids with six synonymous codons (leucine, arginine, serine) only the wobble/third position of the codon was varied from the parent sequence.

#### Degenerate *folA* libraries

The *folA* gene from *E. coli* was manually recoded for reconstruction with degenerate Ultramer^TM^ oligos (IDT). Four oligos were needed to synthesize the full degenerate gene, with junctions designed at methionine or tryptophan codons. Libraries were constructed using scarless assembly reactions to insert oligos into plasmid backbones (BbsI-HF-v2, 1X T4 DNA ligase buffer, T4 DNA ligase; NEB). For amino acids with six synonymous codons (leucine, arginine, serine) only the wobble/third position of the codon was varied from the parent sequence. Six separate sub-libraries of approximately 10,000 variants were created by bottlenecking the larger library for expression level screening (See Materials and methods).

#### Degenerate anti-HER2 VHH libraries

Two regions of an anti-HER2 VHH sequence described in the literature [52] were chosen for investigation as degenerate libraries (codons 2-46, 72-122). To access all possible codons for amino acids with six synonymous codons (leucine, arginine, serine), sub-libraries were constructed using oligos containing either TTR or CTN codons (leucine), TCN or AGY (serine), and CGN or AGR (arginine). These sub-libraries were mixed prior to bacterial transformation. Three versions of the libraries were constructed, a library with only 5’ gene segment degeneracy, a library with only 3’ segment degeneracy and a library with both codon tiles altered. Libraries were assembled similarly to GFP libraries. From each of the three tile library types, approximately 10,000 variants were mixed together to form the final VHH anti-HER2 VHH library.

#### Discrete mCherry and anti-SARS-CoV-2 VHH Strains

Backbone fragments were constructed similarly to GFP library backbones. Discrete strains were assembled using the HiFi DNA Assembly kit (NEB, cat #E2621) from IDT gBlocks. Reactions were assembled with 15 fmol of each backbone fragment and 75 fmol of insert in 20 µL. Reactions were incubated at 50 °C for 60 minutes.

### Library bottlenecking and strain storage

Prepared DNA libraries and discrete plasmids were transformed by electroporation (Bio-Rad MicroPulser) into SoluPro^TM^ *E. coli* B Strain. Cells were allowed to recover in 1 mL SOC medium for 60 min at 30 °C with 250 rpm shaking. For libraries, serial 2-fold dilutions of the recovery outgrowths were plated on Luria broth (LB) agar with 50 µg/mL kanamycin (Teknova) and grown overnight at 37 °C. Transformation efficiency was calculated, and plates with estimated colony numbers closest to our desired diversity were harvested by scraping into LB with 50 µg/mL kanamycin. For discrete strains, single colonies were picked into LB with 50 µg/mL kanamycin and grown overnight at 37 °C with 250 rpm shaking. For discrete strains and libraries, an equal volume of 60 % sterile glycerol was added and 1 mL aliquots were stored frozen at −80 °C.

In the case of mCherry and anti-SARS-CoV-2 VHH discrete strains, plasmid DNA was chemically transformed into SoluPro^TM^ *E. coli* B Strain. Cells were allowed to recover in 1 mL SOC medium for 90 min at 30 °C with 250 rpm shaking. Recovery outgrowths were seeded into LB with 50 µg/mL kanamycin and grown for 42-46 hours. Cultures were stored at −80 °C in 20 % glycerol. Glycerol stocks were plated on LB agar with 50 µg/mL kanamycin (Teknova) and grown overnight at 37 °C. Single colonies were picked into LB with 50 µg/mL kanamycin and grown overnight at 30 °C with 1000 rpm shaking. Cultures were sequence verified. Cultures were stored at −80 °C in 20 % glycerol.

### Protein expression in *E. coli*

Glycerol stocks of bottlenecked libraries or discrete strains in SoluPro^TM^ *E. coli* B strain were diluted into induction base media (IBM, 4.5 g/L Potassium Phosphate monobasic, 13.8 g/L Ammonium Sulfate, 20.5 g/L yeast extract, 20.5 g/L glycerol, 1.95 g/L Citric Acid, adjusted to pH 6.8 with ammonium hydroxide) containing supplements (50 µg/mL Kanamycin, 8 mM Magnesium Sulfate, 1X Korz trace metals).

For GFP library expression, glycerol stocks containing control strains at 0.5 % of cells per strain (3-4 % total) were diluted directly into 25 mL of IBM with supplements and inducers (5 µM arabinose, 5 mM proprionate) and grown for 6 hours at 30 °C with 250 rpm shaking in a baffled flask. Control strains were grown under the same conditions as libraries in 14 mL culture tubes (4 mL volume, 250 rpm shaking) or 96 deep-well plates (1 mL volume, 1000 rpm shaking) depending on experimental need. Cultures were immediately prepared for live-cell Sort-seq assay analysis, or harvested by centrifugation (3000 RCF, 10 min) for downstream biochemical assays.

For GFP and mCherry timecourse expression experiments, seed cultures were created by picking single colonies from strains into 1mL IBM culture and grown overnight at 30 °C, 1000 rpm shaking. Cultures were inoculated with seed into 200 *µ*l IBM with inducers (5 µM arabinose, 5 mM proprionate) in clear 96-well plates at 0.1 OD and grown for 24 hours in a Biotek Synergy H1 plate reader at 30 °C. Area under the curve RFU measurements normalized by OD were collected. A one-way ANOVA statistical test was applied to data points collected to discern statistically significant trends in the different sequence conditions tested for both mCherry and GFP variants.

For *folA* expression under antibiotic selection, glycerol stocks were diluted into 50 mL IBM with supplements and grown overnight at 30 °C with 250 rpm shaking in a baffled flask. Seed cultures were then induced (250 µM arabinose, 1 mM proprionate) and cultured for an additional 2 hours. Induced cultures were then diluted to approximately 50,000 cells per mL in IBM with supplements and inducers (250 µM arabinose, 1 mM proprionate) and grown in 96 deep-well plates with 1 mL volume per well. Control strains were added at a ratio of 5 % total cells. *folA* expression libraries were grown in the presence of sulfamethoxazole (1 *µ*g/*µ*L) (Research Products International, cat #S47000) and a titration of trimethoprim (0, 1, 2, 4, 8, 16, 24, and 32 *µ*g/mL) (Sigma Aldrich) with 0.4 % dimethylsulfoxide. Plates were grown at 30 °C with 1000 rpm shaking (3 mm throw) for 24 hours and cells were harvested by centrifugation (3,000 RCF, 10 min) immediately prior to preparation for sequencing and downstream biochemical analyses.

For VHH expression, glycerol stocks were diluted into LB with 50 µg/mL kanamycin and grown overnight at 37 °C with 250 rpm shaking in a baffled flask or 1000 rpm in 96-well plates. Seed cultures were diluted into IBM with supplements and inducers (250 µM arabinose, 20 mM proprionate) and grown for 22-24 hours at 30 °C with 250 rpm or 1000rpm shaking. Control strains were inoculated from glycerol stocks and grown as for libraries. After 22-24 hours growth, 1 mL aliquots of the induced culture were adjusted to 25 % v/v glycerol and stored at −80 °C before performing downstream biochemical analyses and ACE assays.

### Western blotting

All cell cultures were normalized to OD600= 1 and centrifuged at 1500g for 15 minutes. Cell pellets were resuspended in 200 µL lysis buffer (1X BugBuster^®^ Protein Extraction Reagent, EMD Millipore; 0.025 U/µL Benzonase^®^ Nuclease, EMD Millipore; 1X Halt™ Protease Inhibitor Cocktail, Thermo Scientific; 1 U/µL rLysozyme™, EMD Millipore), incubated at room temperature for 30 minutes, and centrifuged at 4000 g for 30 minutes. Supernatant was removed and stored as soluble material. The remaining pellet was resuspended in 200 µL lysis buffer and stored as insoluble material. Laemmli sample buffer (1X final) and dithiothreitol (DTT, 100 mM final) were added to insoluble and soluble fractions and incubated at 70 °C for 20 min. Samples were run on Novex WedgeWell 4-20 % Tris-Glycine Gels (Invitrogen) in Tris-Glycine SDS Running Buffer (225V, 45 min)and transferred to nitrocellulose (iBlot 2, iBlot 2 NC Mini Stacks; Invitrogen) at 20 V for 1 min, 23 V for 4 min, and 25 V for 2 min. Blots were incubated in blocking buffer (3 % BSA in tris-buffered saline plus 0.05 % Tween-20 [TBS-T]) for one hour at room temperature or 4C overnight. Quantification was performed via densitometry using AzureSpot Pro (Azure Biosystems) with rolling-ball background correction.

### GAPDH blots

For all glyceraldehyde phosphate dehydrogenase (GAPDH) quantification, blots were cut in half below the 35 kDA marker. The upper half was then probed with 1:1000 GAPDH Loading Control Monoclonal Antibody, Alexa Flour 647 (MA5-15738-A647, Invitrogen) in blocking buffer (1 h room temp) and imaged using an Azure600 (Cy5 fluorescence channel).

### GFP Western blots

‘Blots were incubated in 1:2000 GFP Polyclonal Antibody (A-11122, ThermoFisher Scientific) in blocking buffer (1 hour, room temp), 1:2500 Goat anti-Rabbit IgG (Heavy Chain), Superclonal™ Recombinant Secondary Antibody, HRP (cat #A27036, ThermoFisher Scientific) in blocking buffer (30 min room temp). SuperSignal PLUS Chemiluminescent substrate (ThermoFisher Scientific) was added and the membrane was incubated at room temperature for 5 minutes and imaged on an Azure300.

### DHFR Western blots

Blots were incubated in 1:5000 VHH against DHFR/*folA* (cat #PNBL047, Creative Biolabs) in blocking buffer for (1 hour room temp), 1:500 Goat anti-alpaca Alexa647 (cat #128-605-230, Jackson ImmunoResearch) in blocking buffer (30 min room temp), and imaged on an Azure600 (Cy5 fluorescence channel).

### Anti-Her2 VHH Western blots

Blots were incubated in 1:2000 Anti-polyhistidine-Alkaline Phosphatase antibody, Mouse monoclonal (cat #A5588, Sigma-Aldrich) in blocking buffer (1 hour room temp). 1-Step™ NBT/BCIP Substrate Solution (ThermoScientific) was added (5 min room temp) and the membrane was washed with deionized water then imaged on an Azure300.

### GFP variant measurements via Sort-seq assay

To generate high-throughput expression measurements of synonymous DNA sequence variants of GFP we applied a FACS based sort-seq protocol similar to Schmitz et al. [16] to three separate tiled degenerate libraries. For staining, aliquots of OD_600_ = 2 from induced cultures were made in 0.7 mL matrix tubes, centrifuged at 3300 g for 3 min, and pelleted cells were washed 3X with 1X PBS + EDTA. Washed cells were then resuspended in 500 µL of 1X PBS + EDTA for sorting. Libraries were sorted on FACS Symphony S6 instruments. Prior to sorting, 40 µL of sample was transferred to a FACS tube containing 1 mL PBS+EDTA with 3 µL propidium iodide. Aggregates, debris, dead cells were excluded using singlets, size, and PI-parent gates, respectively. The GFP-positive population was divided into 6 evenly spaced gates with 400,000 events collected per gate in 3 replicates. Sorted cells were then centrifuged for 10 minutes at 3800 g, washed once with 450 µL of DI water, resuspended, and centrifugred for 10 minutes at 3800 g. Supernatant was aspirated and samples were processed for DNA extraction and NGS analysis.

The three tile libraries were screened in isolation, with tile 1 screened on a separate day than tile 2 and tile 3. Fluorescence values of the libraries were normalized based on concurrently measured control variants (fig. S5,fig. S6). The normalized expression score values allowed for combination of tiles into a single dataset (fig. 2B). Sequencing data from the Sort-seq method was used to generate expression score values and subjected to quality thresholding (See Materials and methods). Expression score values were validated using a subset of 24 variants per tile measured via Plate Reader, Cytometer and soluble protein levels via Western Blot (fig. S7), with high correlation between all metrics (fig. S8).

### *FolA* expression measurements via antibiotic selection

Cells expressing synonymous DNA sequence variants of *folA* prepared and grown as described. Plasmid DNA was extracted and PCR amplicons were generated and sequenced in the described manner. Sequence counts were used to generate weighted expression scores as described.

### VHH variant measurements via Activity-specific Cell-Enrichment (ACE) assay

We applied a modified version of the previously described ACE assay to generate high-throughput protein level measurements of anti-Her2 and anti-SARS-CoV-2 VHH synonymous DNA sequence variants expressed in SoluPro^TM^ *E. coli* B strain [42].

#### Cell Preparation

High-throughput screening was performed on VHH codon libraries intracellularly stained for functional antigen binding. An OD_600_ = 2 of thawed glycerol stocks from induced cultures were transferred to 0.7 mL matrix tubes and centrifuged at 3300 g for 3 min. The resulting pelleted cells were washed three times with PBS + 1 mM EDTA and thoroughly resuspended in 250 µL of 32 mM phosphate buffer (Na_3_HPO_4_) by pipetting. Cells were fixed by the addition of 250 µL 32 mM phosphate buffer with 1.3 % paraformaldehyde and 0.04 % glutaraldehyde. After 40 min incubation on ice, cells were washed three times with PBS, resuspended in lysozyme buffer (20 mM Tris, 50 mM glucose, 10 mM EDTA, 5 µg/mL lysozyme) and incubated for 8 min on ice. Fixed and lysozyme-treated cells were equilibrated by washing 3x in stain buffer.

#### Staining

After lysozyme treatment and equilibration, the Her2 VHH library was resuspended in 500 µL Triton X-100 based stain buffer (AlphaLISA immunoassay assay buffer from Perkin Elmer; 25 mM HEPES, 0.1 % casein, 1 mg/mL dextran-500, 0.5 % Triton X-100, and 0.05 % kathon) with 50 nM human HER2:AF647 (Acro Biosystems) and 30 nM anti-VHH probe (MonoRab anti-Camelid VHH [iFluor 488], GenScript cat #A01862). The SARS-CoV-2 VHH strains were resuspended in saponin based stain buffer (1x PBS, 1mM EDTA, 1 % heat inactivated fetal bovine serum, and 0.1 % saponin) with 75 nM SARS-CoV-2 delta RBD:AF647 (Acro Biosytems) and 25 nM anti-VHH probe. Samples were incubated with probe overnight (16 hours) with end-to-end rotation at 4 °C protected from light. After incubation, cells were pelleted, washed 3x with PBS, and then resuspended in 500 µL PBS.

#### Analysis and Sorting

Immediately prior to screening, 50 µL prepped sample was transferred to a flow tube containing 1 mL PBS + 3 µL propidium iodide. Aggregates, debris, and impermeable cells were were removed with singlets, size, and PI^+^ parent gating. The SARS-CoV-2 VHH strains were screened on a Sony ID7000 spectral analyzer. Anti-Her2 VHH libraries were sorted using FACSymphony S6 (BD Biosciences) instruments. Collection gates were drawn to evenly fraction the log range of probe signal (fig. S3). The pooled VHH library tiles were sorted twice sequentially on the same instrument. The collected events were processed independently as technical replicates.

### Amplicon generation for NGS

#### Post-selection *folA* amplification

DNA was extracted from bacterial cultures grown under selection conditions by miniprep (Qiagen, Qiaprep 96 cat #27291 or Qiaprep cat #27106). The folA variable region was amplified by PCR (Phusion^TM^, ThermoFisher cat #F530 or Q5^®^, NEB cat #M0492) with 500 nM primer. See supplemental oligo file for oligo sequences and PCR conditions. PCR reactions were then purified using ExoSAP-IT PCR Product Cleanup Reagent (ThermoFisher), quantified by Qubit fluorometer (Invitrogen), normalized, and pooled. Pool size was verified via Tapestation 1000 HS and sequenced.

#### Sort-seq GFP and ACE assay anti-HER2 VHH amplification

Cell material from various gates was collected in a diluted PBS mixture (VWR), in 96 well plates. Post-sort samples were spun down at 3,800 g and tube volume was normalized to 20 µl. Amplicons for sequencing were generated via PCR, using collected cell material directly as template with 500 nM primer concentration, Q5 2x master mix (NEB) and 20 µl of sorted cell material input suspended in diluted PBS (VWR). See supplemental oligo file for oligo sequences and PCR conditions. PCR reactions were then purified using using ExoSAP-IT PCR Product Cleanup Reagent (ThermoFisher), quantified by Qubit fluorometer (Invitrogen), normalized, and pooled. Pool size was verified via Tapestation 1000 HS and sequenced.

### Sequencing

Amplicons were prepared for sequencing using Takara ThruPLEX^®^ DNA-Seq Kit (Takara Bio, cat #R400674), which included unique dual indexing. To ensure a minimum of 50 bp read overlap, libraries with insert sizes of 250 bp or greater were sequenced using 2×300 paired-end reads on an Illumina MiSeq using a MiSeq Reagent Kit v3 (Illumina Inc, MS-102-3003). Libraries with insert sizes of less than 250 bp were sequenced using 2×150 paired-end reads. These were sequenced on either an Illumina MiSeq or NextSeq, depending on the read depth required for each run. Each run was sequenced with a 20 % PhiX spike-in for diversity.

### Sequence processing

In order to convert sequence counts from sorting and selection procedures, the following processing and quality control steps were performed:

1. Adapter sequences were removed using the CutAdapt tool. [65]
2. Sequencing reads were merged using Fastp [66] with the maximum overlap set according to the amplicon size and the read length used in each experiment.
3. Primer sequences were removed from both ends of merged reads using the CutAdapt tool, [65] and reads without the primer sequences were discarded.
4. Raw counts and PPM counts were calculated for each variant using custom Python scripts.

### High-throughput dataset sequencing count based expression score calculation

To compute expression scores for the three protein datasets, we performed the following processing and quality control steps:

1. Variants were filtered to remove DNA sequences that did not translate to the correct sequence of the target protein region.
2. The total count across all gates or antibiotic concentrations was computed for each replicate. Variants with fewer than 10 total counts in either replicate were discarded.
3. For each gate or antibiotic concentration, the variant counts were normalized by the total count (in millions).
4. The expression score for each variant in the Sort-seq and ACE experiments was computed using a weighted average using the log-transformed geometric mean fluorescence intensity within each gate (log10 MFIgate):

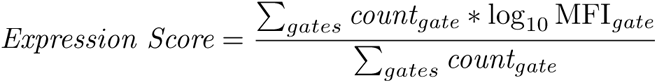 The following weighted average was used for the antibiotic selection experiment:

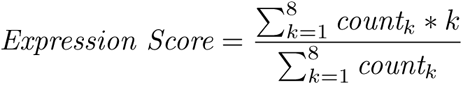

Where *k* is the integer rank of the antibiotic concentration (i.e. *k* = 1 represents the lowest concentration whereas *k* = 8 represents the highest concentration).
5. Expression scores were averaged across independent FACS sorts or replicates of antibiotic selection.
6. Specifically for the GFP data, GFP tile 1 was measured separately from GFP tiles 2 and 3. In order to reconcile batch variability in measurements, the following normalization procedure was performed. During the FACS sort, 10 sequences per tile were included as spike-in controls. The fluorescence of all 30 spike-in variants was measured alongside the tiled libraries using an FACS Symphony S6. Linear regression was performed on the log transformed mean fluorescent intensity (log-MFI) of the spike-ins to determine a scaling function that could translate log-MFI values from the tile 1 distribution to the tile 2/3 distributions (fig. S5). This function was applied to the expression scores in tile 1, resulting in a consistent expression score for the parent GFP sequence (fig. S6)

### Protein2DNA dataset construction

To construct the Protein2DNA dataset, we downloaded protein and DNA records from RefSeq as well as taxonomic identifiers from the NCBI Taxonomy database on July 1, 2021. We considered all taxonomic kingdoms when downloading these records. Next, we filtered the genomic records according to their metadata. Specifically, we included only genomes with a status of “reference genome” or “representative genome”. We also filtered out incomplete genomes by using the “Complete Genome” or “Full” tags in the record metadata. Each record was scanned for coordinate alignment between the DNA and its amino acid translation, dropping those records with inconsistent protein-DNA mapping. For genes, records labeled as “pseudo” were excluded and only those with a “cds” or “translation feature” tag were considered. Then, we ensured that only records with a valid stop codon were included. For each corresponding protein sequence, we ensured that the sequence was not truncated (that is, the protein sequence starts with an “M” and ends with a “*” or stop symbol). Finally, only sequences with canonical DNA bases or amino acids were included in the dataset.

### Processing of Protein2DNA dataset entries

In order to train our language models, we created a dictionary mapping relevant characters or words from the Protein2DNA dataset to unique *tokens*. Briefly, we assigned unique tokens to (1) each of the 20 amino acids as well as the stop symbol; (2) each of the 64 codons; (3) each of the taxonomic ranks (Kingdom, Phylum, Class, Order, Family, Genus, Species, Strain, and Genetic Code); and (4) each of the 10 numeric digits (0-9) to represent taxonomic identifiers. In this way, each entry of our Protein2DNA dataset is converted to a language model-compatible representation by tokenizing each of the words comprising its taxonomy, amino acid sequence, and DNA sequence. Specifically, the order of the tokenized words for a Protein2DNA entry whose corresponding CDS has *N* codons is: [<KINGDOM> <KINGDOM number> <PHYLUM> <PHYLUM number> <CLASS> <CLASS number> <ORDER> <ORDER number> <FAMILY> <FAMILY number> <GENUS> <GENUS number> <SPECIES> <SPECIES number> <STRAIN> <STRAIN number> <1ST amino acid> <2ND amino acid> … <(N-1)th amino acid> <STOP symbol>] for the input text, and `[<1ST codon> <2ND codon>… <NTH codon>]` for the label.

### Training of CO-T5 model

Our CO-T5 architecture is based on the `T5ForConditionalGeneration` model [43] and its PyTorch implementation within the HuggingFace framework [67]. The model contains 12 attention layers, the hidden layer size is 768 and the intermediate layer size is 1152. Model training was performed in a supervised manner, whereby the model is shown a tokenized input text (taxonomy + amino acid sequence) and the corresponding tokenized label text (CDS sequence) as described above. We used a learning rate of 4 10*^−^*^4^ with 1000 warm-up steps and no weight decay. The final CO-T5 model was trained for 84 epochs, which corresponded to the point where both the training and validation loss had converged while avoiding overfitting.

### Training of CO-BERTa model

Our CO-BERTa architecture is based on the `RobertaForMaskedLM` model [46] and its PyTorch implementation within the HuggingFace framework [67]. The model contains 12 attention heads and 16 hidden layers. The hidden layer size is 768 and the intermediate layer size is 3072. Model training in a self-supervised manner following a dynamic masking procedure with a special `<MASK>` token. For masking, we used the `DataCollatorForLanguageModeling` class from the Hugging Face framework to randomly mask codon tokens with a probability of 20%. Entries from the Protein2DNA datasets were processed in the same way as for the CO-T5 model. Training was performed with the LAMB optimizer [68] with learning rate of 10*^−^*^5^, linear rate decay with weight of 0.01, 1000 steps of warm-up, a gradient clamp value of 1, and dropout probability of 20%. The model was trained for 100 epochs, which corresponded to the point where both the training and validation loss had converged while avoiding overfitting.

### Finetuning of CO-BERTa model

We fine-tuned our pre-trained CO-BERTa model by adding a dense hidden layer with 768 nodes followed by a projection layer with the a single output neuron (regressor). All layers remained unfrozen to update all model parameters during training. Training was performed with the AdamW optimizer [69], with a learning rate of 10*^−^*^5^, a weight decay of 0.01, a dropout probability of 0.2, a linear learning rate decay with 1000 warm up steps, and mean-squared error (MSE) as the loss function.

### Synonymous DNA Expression Datasets

The three datasets (GFP, anti-HER2 VHH, and folA) outlined above were used to finetune the pre-trained MLM CO-BERTa model as a sequence-to-expression predictor. Each dataset was first filtered to include sequences with at least 10 read counts, resulting in the following dataset sizes: GFP=119,703 sequences, anti-HER2 VHH=17,146 sequences, and folA=17,319 sequences. These were then used to fine-tune the pre-trained model in three different ways: (1) using the GFP dataset alone, (2) using the anti-HER2 VHH dataset alone, and (3) using sequences from all three datasets. Since the GFP dataset was significantly larger than the other two, we randomly sampled 18,000 GFP sequences for fine-tuning the model in this last case. In each case, 90% of all sequences were used for fine-tuning and the remaining 10% were held out as validation and test sets (5% of the dataset, respectively).

### Sequence Naturalness score

We define the naturalness *n_s_* of a sequence as the inverse of its pseudo-perplexity. Recall that, for a sequence *S* with *N* tokens, the pseudo-likelihood that a model with parameters Θ assigns to this sequence is given by:

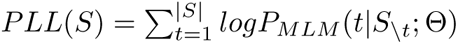

The pseudo-perplexity is obtained by first normalizing the pseudo-likelihood by the sequence length and then applying the negative exponentiation function:

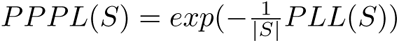

Thus, the sequence Naturalness score is:

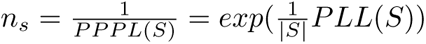

Naturalness scores were computed using the pre-trained CO-T5 model described above.

### Sequence metrics (CAI, GC%, and %MinMax)

For each DNA sequence generated by our pre-trained CO-T5 model, we computed three metrics as described before: Codon Adaptation Index (CAI) [44], GC% content, and %MinMax [26].

Briefly, the CAI of a DNA coding sequence with N codons is the geometric mean of the frequencies of its codons in the original source genome.

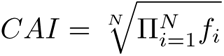

GC% is the fraction of nucleotides in the DNA coding sequence that are either Guanine (G) or Cytosine (C). The algorithm for computing %MinMax has been described in detail elsewhere [26]. In our implementation, we used a codon window size of 18.

### Generation and model scoring of *in silico* degenerate codon libraries

Using our fine-tuned CO-BERTa models, we scored 10 million random synonymous DNA coding variants *in silico* for each mCherry and anti-COVID VHH. We restricted the insertion of random synonymous codons to the same tiles used in the GFP and anti-HER2 VHH libraries, to ensure that we maintained similarity to the datasets used during fine-tuning. For the mCherry library, we scored sequences with both the GFP and the ALL models, whereas for the anti-COVID VHH library we used the anti-HER2 and the ALL models. We selected the top 10 best and bottom 10 worst scored sequences for each library as scored by each of the two corresponding models. Additionally, we randomly sampled codons to generate random 10 random CDSs for downstream experimental validation.

### XGBoost baseline model training

We trained baseline XGBoost models using two different approaches. First, using our pre-trained CO-T5 model, we generated embeddings for all sequences in the Synonymous DNA Expression Datasets (GFP, folA, and anti-HER2 VHH). We then trained three individual XGBoost models with each dataset by using a random split of 90% for training and 5% for validation and test holdout each. This first type of models were dubbed **XGBoost CO-T5**. For the second approach, we trained the same three XGBoost models except that each sequence was converted to a one-hot encoding by mapping each of the 64 codons to a unique value from 1 through 64. These models are dubbed **XGBoost 1HE**. For all models, we used the **xgboost** package for Python (version 1.7.2) and the **xgboost.train** function with the following hyperparameters: *nboosts = 100*, *eta = 0.1*, *booster = ‘gbtree’*, *objective = ‘reg:squarederror’*.

## Supplementary Information

**Fig S1.**
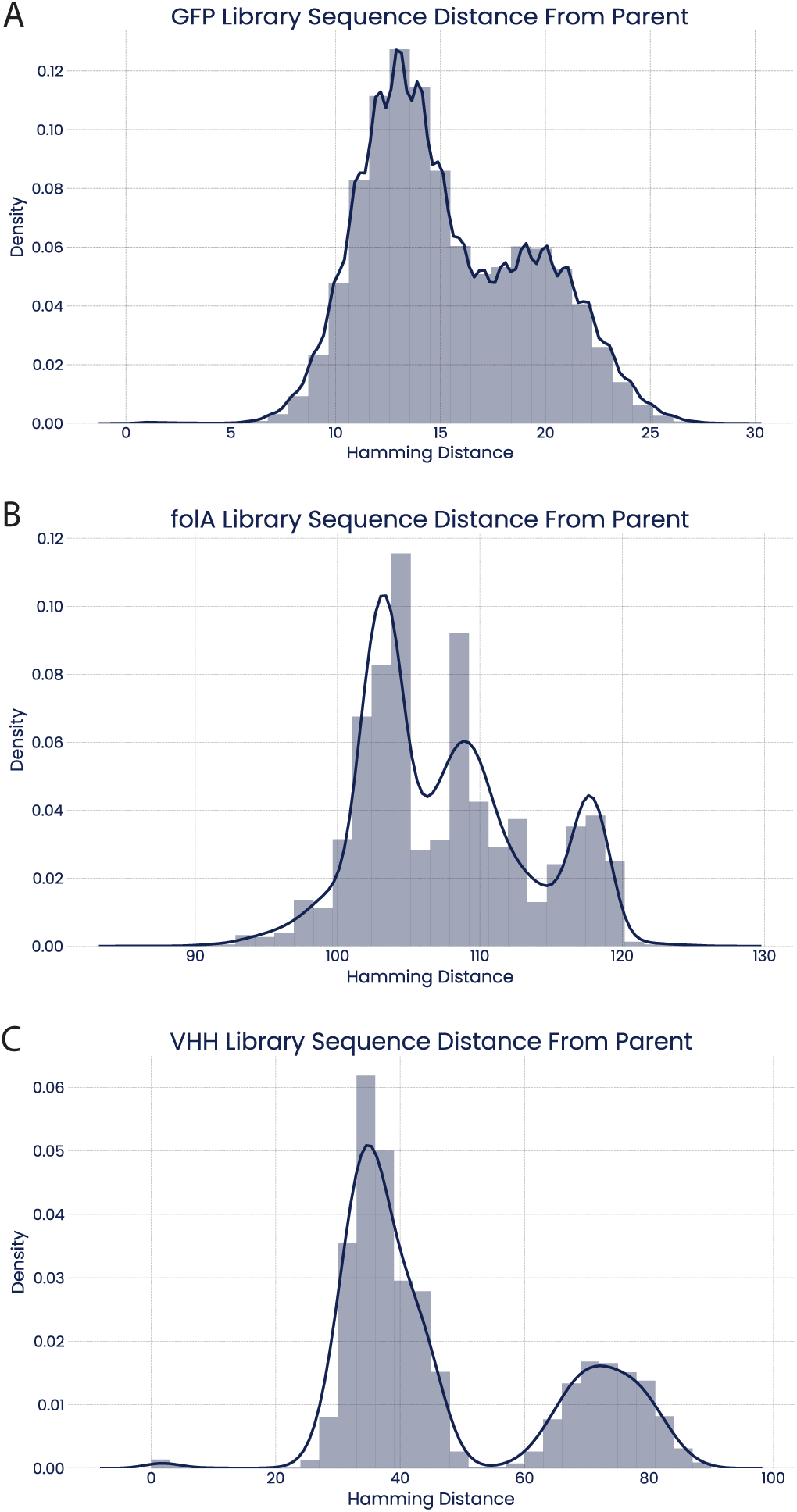
DNA sequence diversity in expression libraries. DNA Hamming distance from parent sequence for all three synonymous protein libraries. **(A), (B), (C)** Nucleotide level Hamming distance from parent for all three protein libraries used in the study. Plots show diverse DNA sequence space sampled within all libraries.

**Fig S2.**
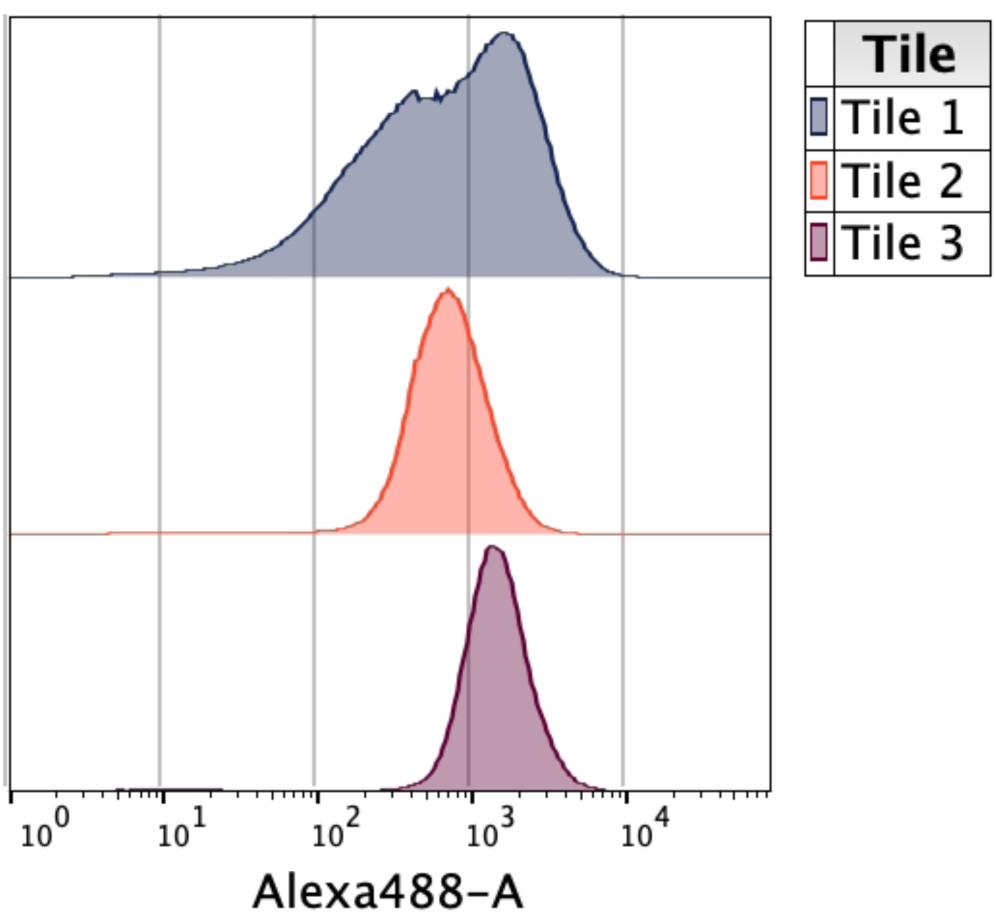
Fluorescence distribution of GFP tile libraries. Debris, aggregates, and dead cells were excluded by parent gating prior to plotting the fluorescence signal for each library.

**Fig S3.**
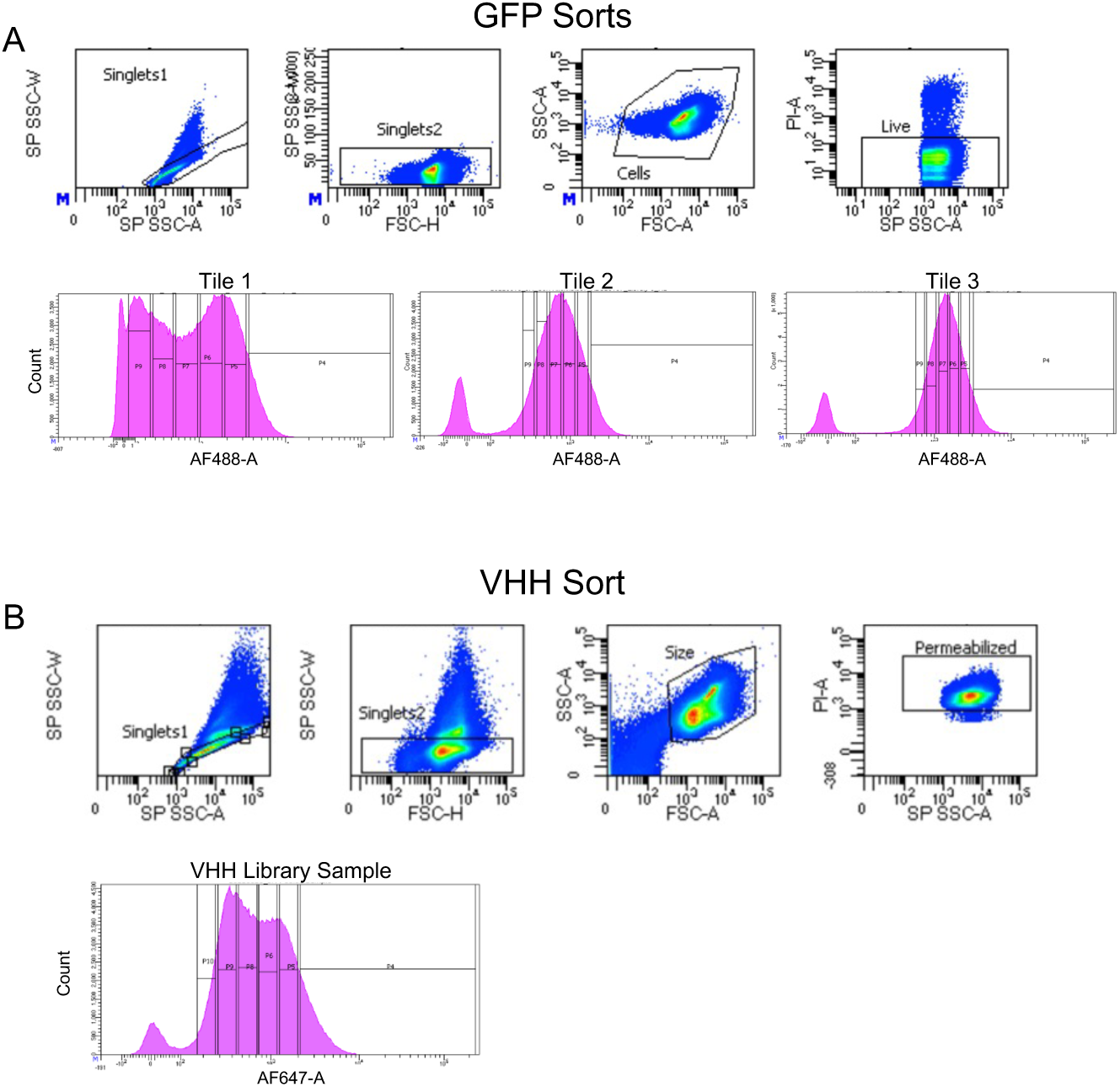
Gating schemes for library sorting. **(A)** Representative parent gating for GFP library sorts is shown. The two singlets gates were drawn to exclude cellular aggregates regions previously identified by dual fluorescence of GFP and mCherry reporter strains. Propidium iodide (PI) was used to exclude dead cells. Collection gates are shown for each tile. **(B)** Parent gating for the VHH library was similar as described above except PI was used to exclude non-permeabilized cells. For both GFP and VHH libraries, six collection gates were drawn to sample across the range of fluorescence distribution of cells expressing functional protein.

**Fig S4.**
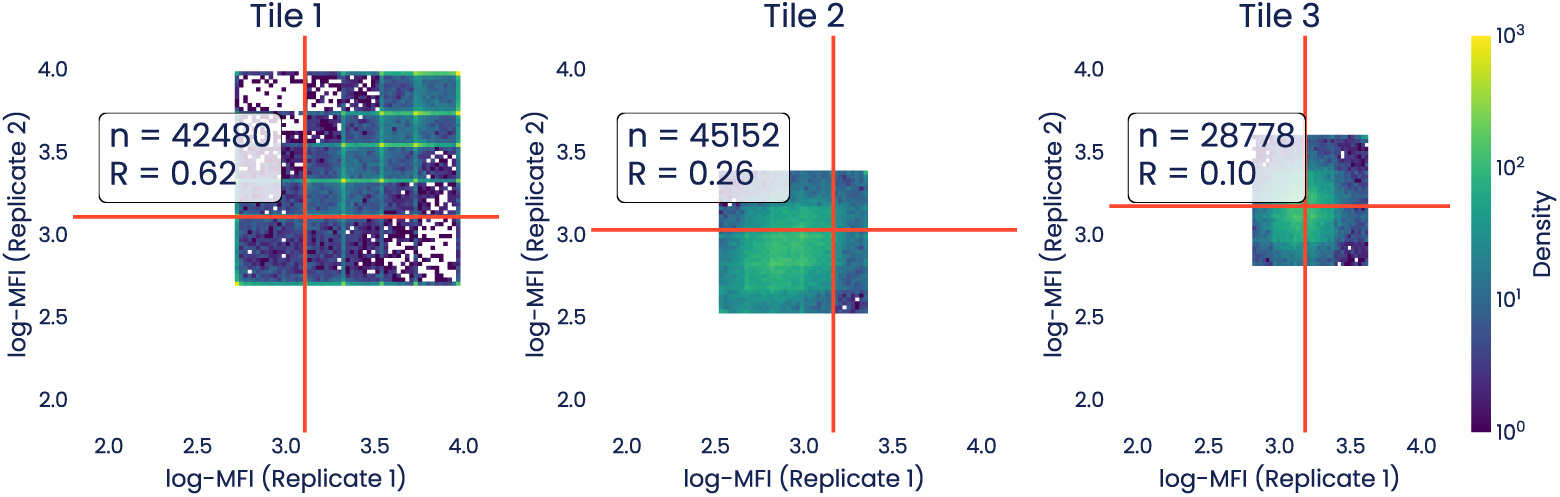
Density heatmap of expression scores between replicate sorts for the three GFP library tiles. Red lines indicate the parent GFP log-MFI. Pearson R correlation coefficients and number of datapoints passing quality control filters are shown for each tile.

**Fig S5.**
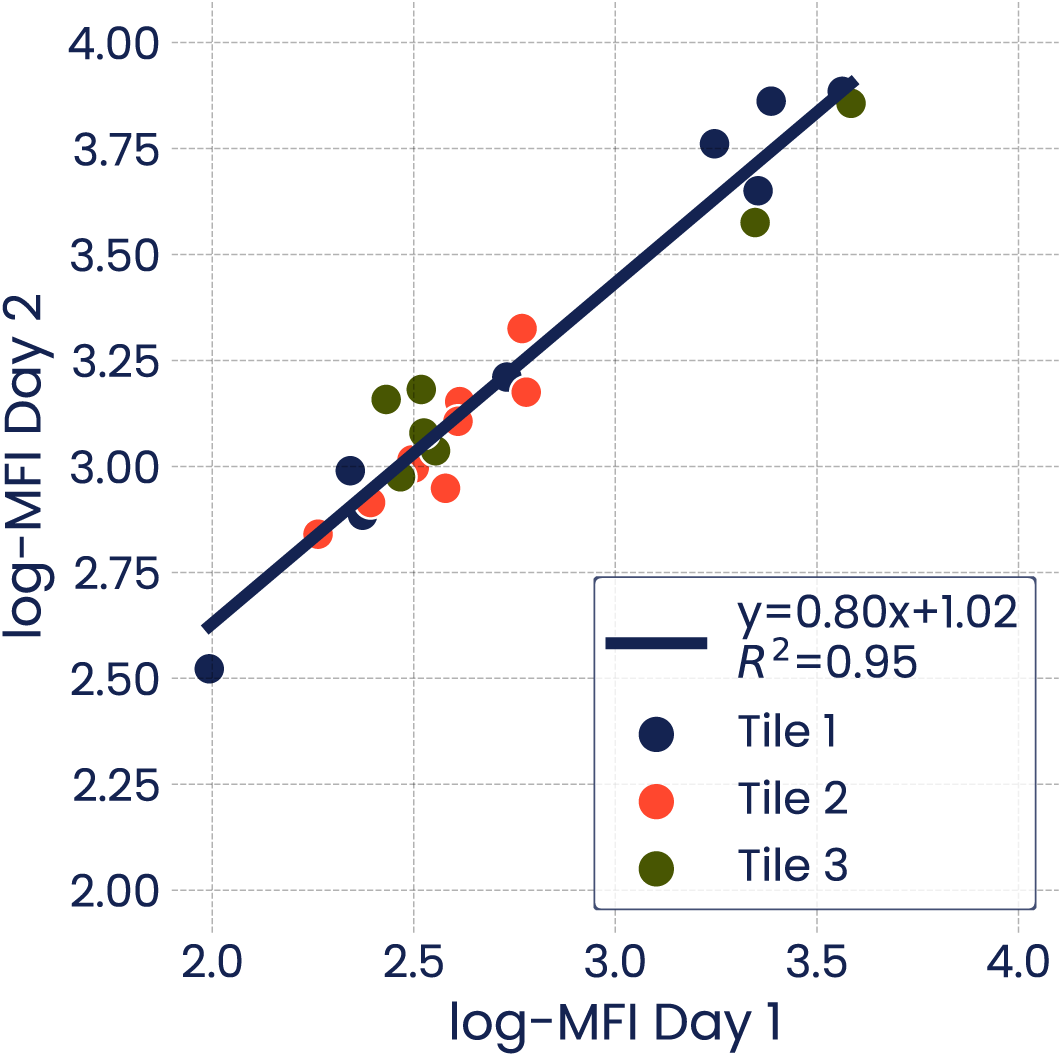
Scatterplot showing log-MFI of control sequences. Measurements on Day 1 (tile 1) or Day 2 (tiles 2 and 3) are shown. WT refers to the parental GFP sequence from which the libraries were derived.

**Fig S6.**
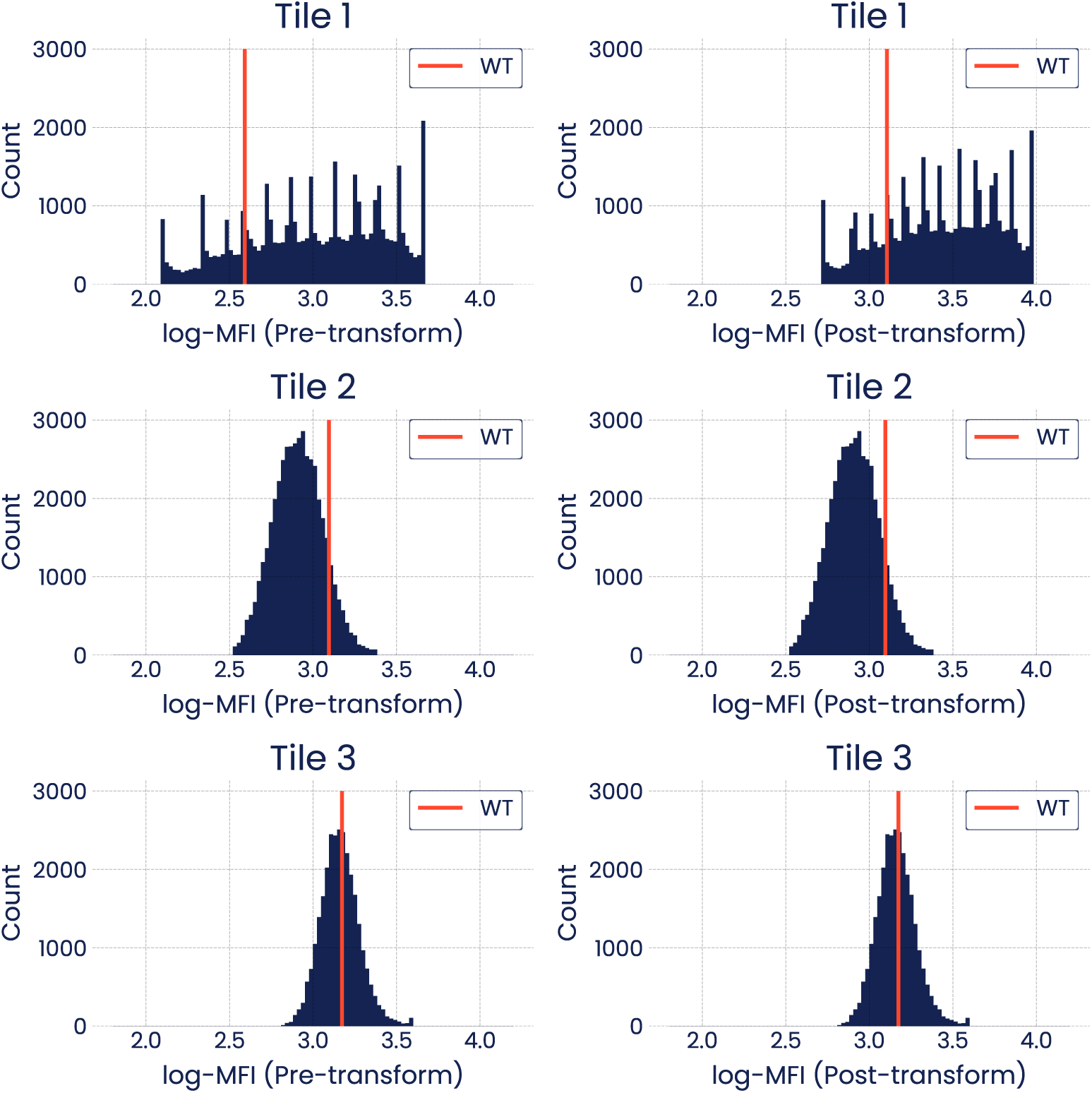
Log-MFI expression score histograms for each tile before and after normalization. Red vertical line indicates parent GFP Sequence log-MFI. WT refers to the parental GFP sequence from which the libraries were derived.

**Fig S7.**
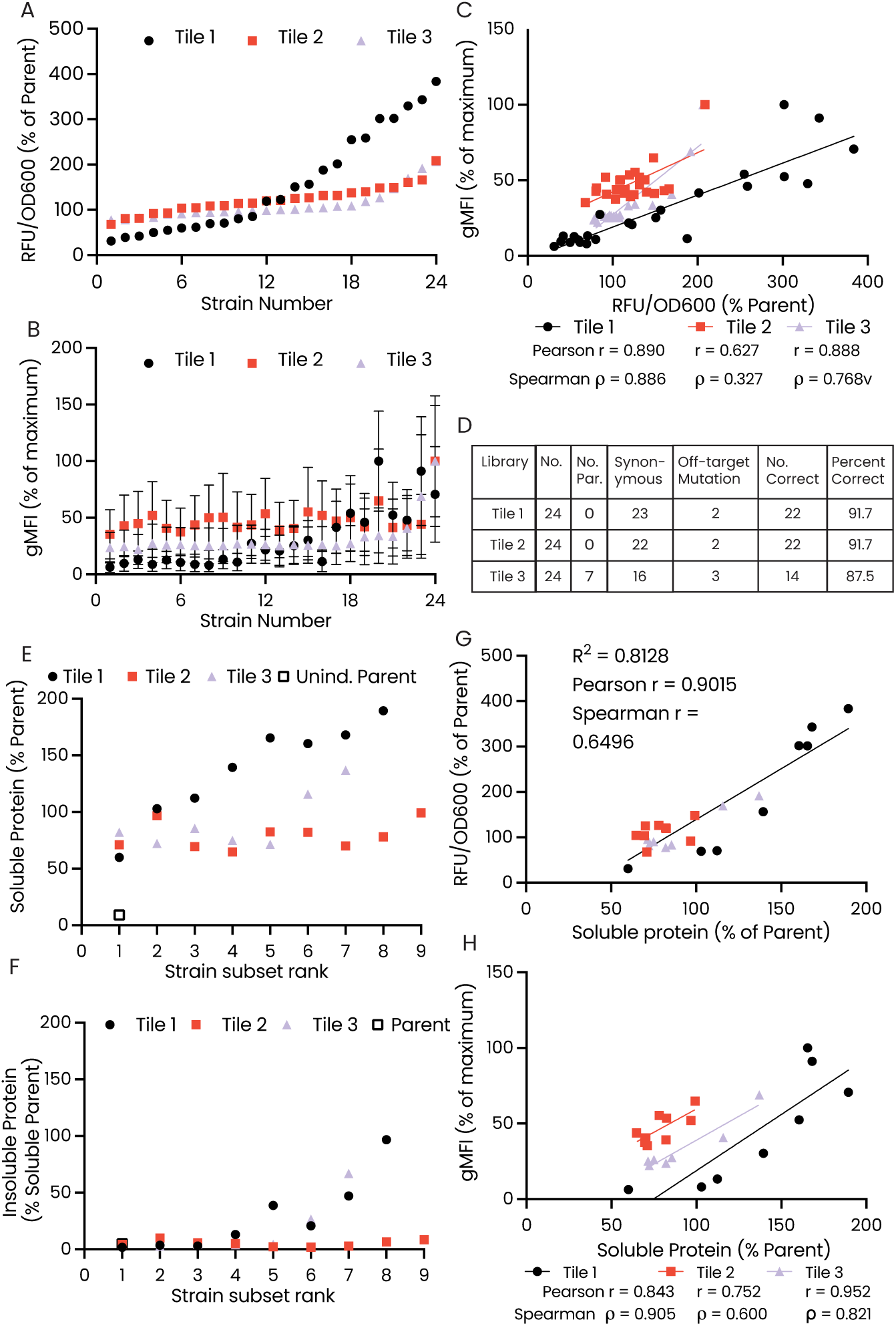
Validation of GFP degenerate codon library control variants. **(A)** Plate reader RFU/OD600 values from 24 variants from the three GFP tile libraries. **(B)** Fluorescent cytometry readings of the 24 variants from each tile. Error bars are standard deviation of recorded events. **(C)** Correlations of Plate Reader and Cytometer fluorescent measurements. **(D)** Table of sequencing characterization of 24 colonies from each tile. Minimal levels of non-GFP coding variants in sampled variants. **(E)** Soluble protein levels as measured by Western Blot of a subset of variants per tile. Values are reported as a percentage of parent GFP sequence levels. **(F)** Insoluble protein levels of variants from panel E as measured by Western Blot. Values reported as a percentage of parent sequence soluble protein level. **(G)** Plate Reader fluorescent values correlated with soluble protein levels as measure by western blot. Plot shows variants from each tile appearing in panel E. **(H)** Correlation of Western Blot soluble protein to geometric mean fluorescence of variants from each tile.

**Fig S8.**
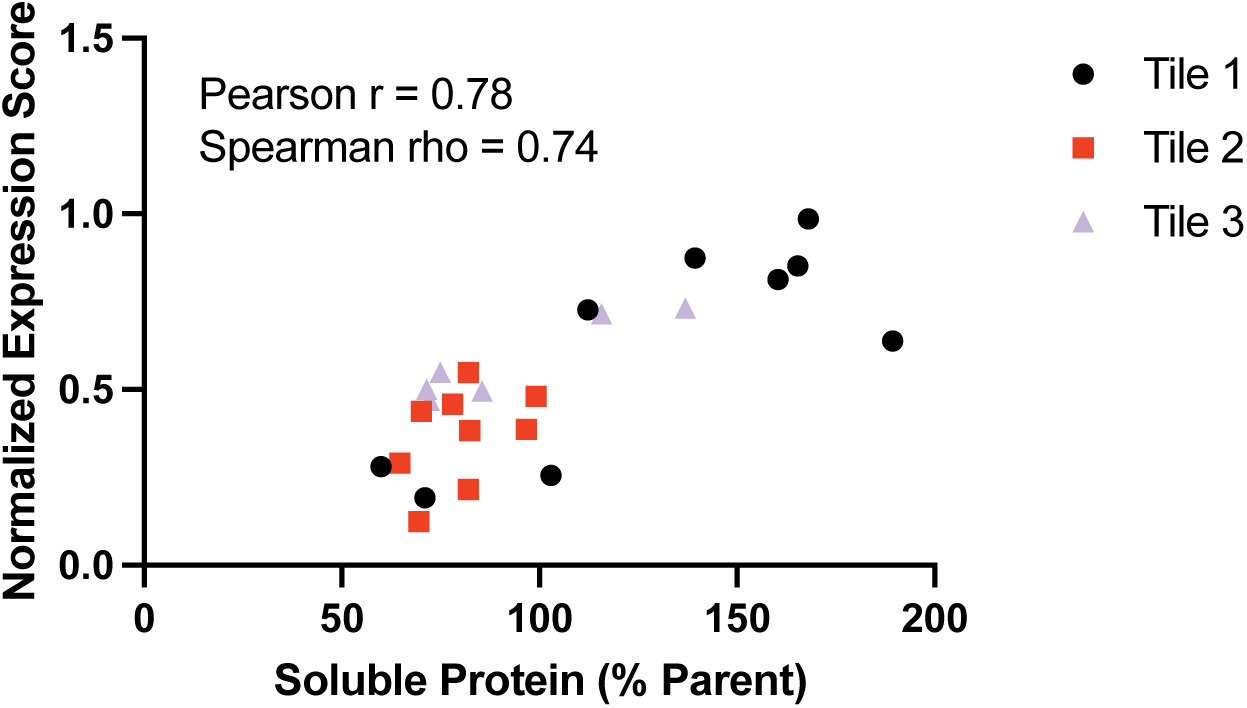
Soluble protein level of GFP variants measured via western blot vs Normalized Expression Score. Plot shows Sort-seq derived expression scores and soluble protein levels as measured via Western Blot of representative GFP variants from S7. Correlation is observed between the two measurement types across variants from all three tiles.

**Fig S9.**
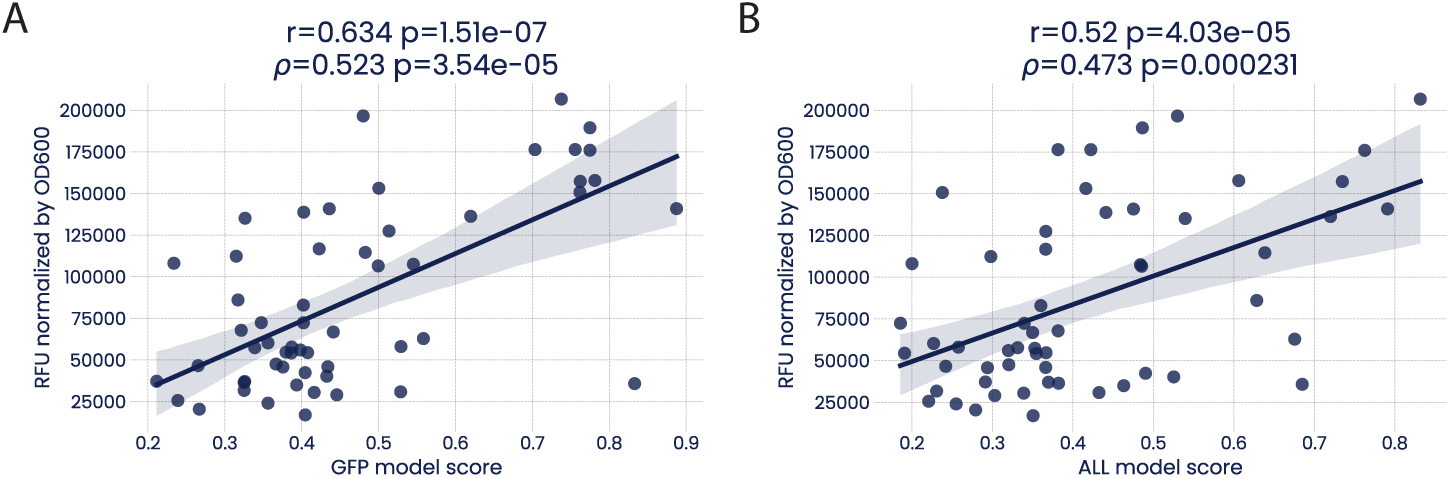
Fine-tuned expression level predictions on CO-T5 generated and commercial GFP Sequences. **(A)** Correlation of GFP model expression level predictions for GFP sequences from fig. 1E and plate reader fluorescence measurements. **(B)** Correlation of ALL model expression level predictions for GFP sequences from fig. 1E and plate reader fluorescence measurements.

**Fig S10.**
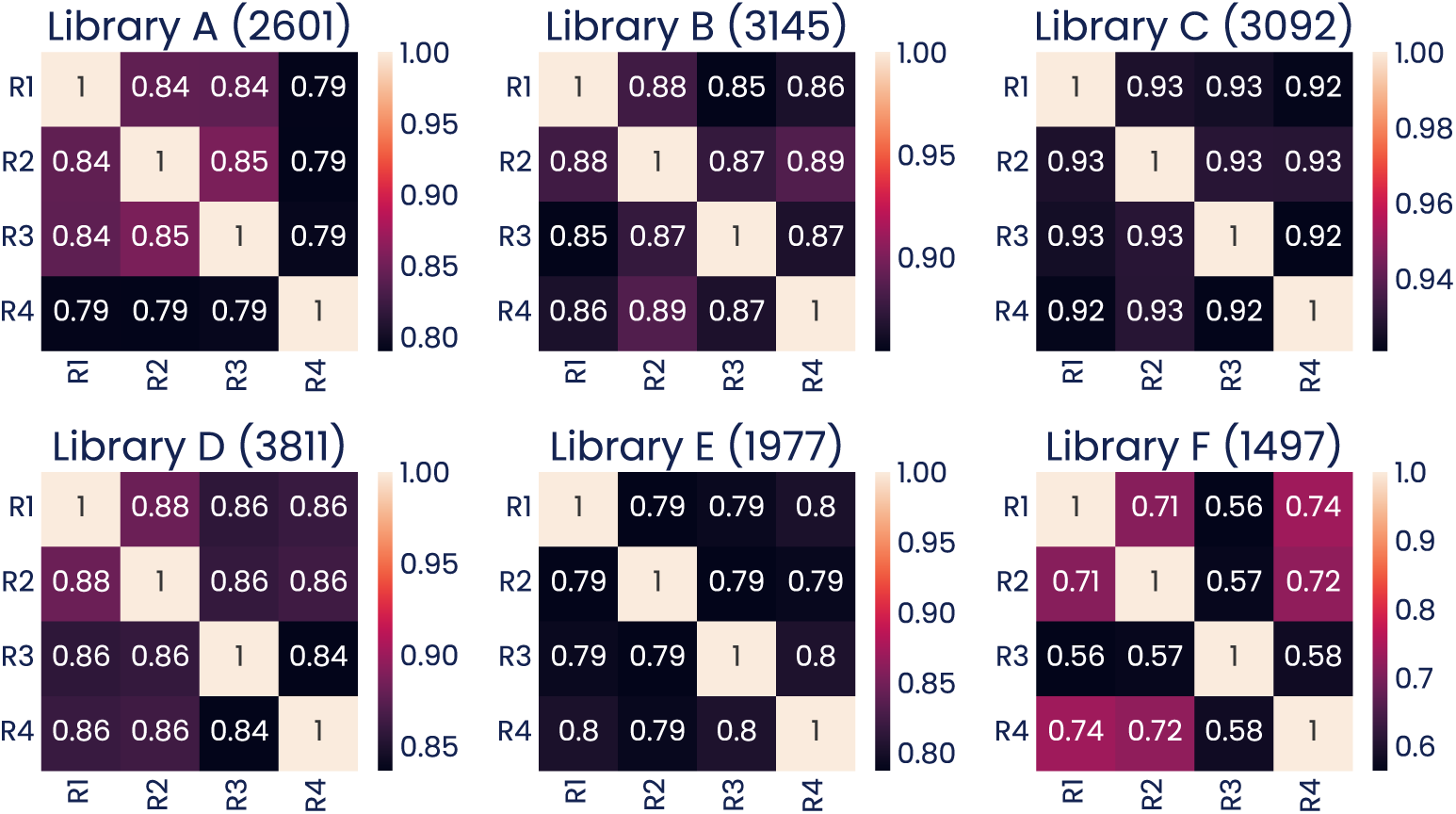
Heatmap of all-vs-all expression score Pearson R correlations for *folA* sub-libraries. Correlations between the four replicate screens from each of the six *folA* sub-libraries. Numbers in parentheses indicate number of variants in each library that passed all quality control filters.

**Fig S11.**
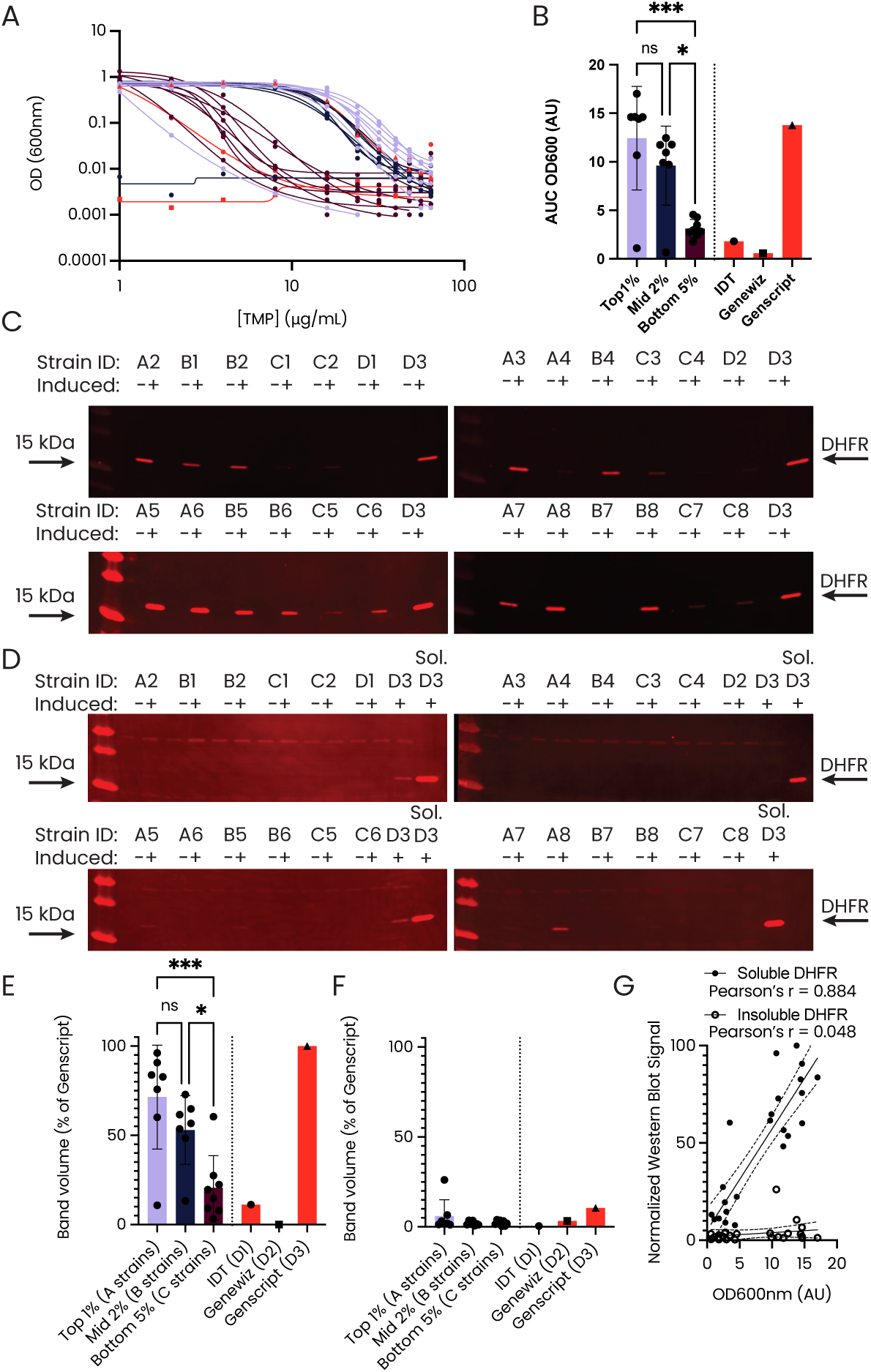
Validation of DHFR drug selection score. **(A)** Dose-response curves of synonymous *folA* codon variants from the top 1 % (lavender), middle 2 % (black), and bottom 5 % (maroon) of score range from drug selection. Commercially optimized variants are shown in orange. Cultures were grown for 24 hours after expression induction in the presence of 1 µg/mL SMX and the indicated TMP concentration. **(B)** Area under the curve (AUC) of data in (A) from lines fit using GraphPad Prism. **(C)** Anti-DHFR Western blots on soluble protein fraction of cell lysates from *folA* strains shown in (A) and (B). **(D)** Anti-DHFR Western blots on insoluble protein fraction of cell lysates from *folA* strains shown in (A) and (B). **(E)** Quantification of data in (C), normalized to signal from D3. **(F)** Quantification of data in (D). G) Correlation of soluble and insoluble DHFR protein expression with AUC data in (B), normalized to signal from soluble fraction D3. Statistics and AUC calculations were performed in GraphPad Prism. Two-way ANOVA tests were performed in panels B, E, F between strain groups A, B, and C where *** = p*<*0.001, ∗ = *p<*0.05*, ns* = *p>*0.05.

**Fig S12.**
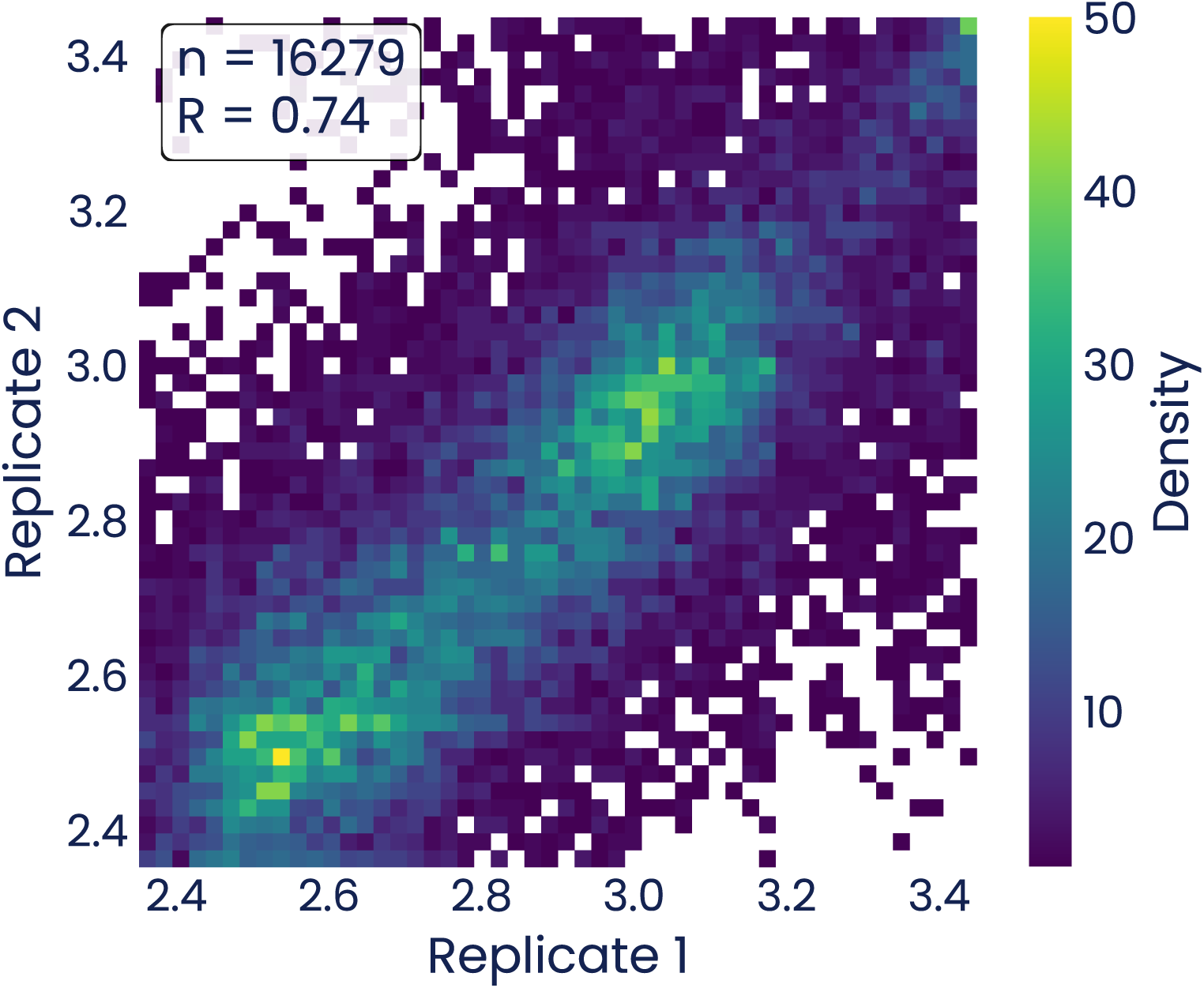
Density heatmap of ACE expression scores between replicate sorts for the VHH library. Pearson R correlation coefficient and number of datapoints passing quality control filters are shown in the legend.

**Fig S13.**
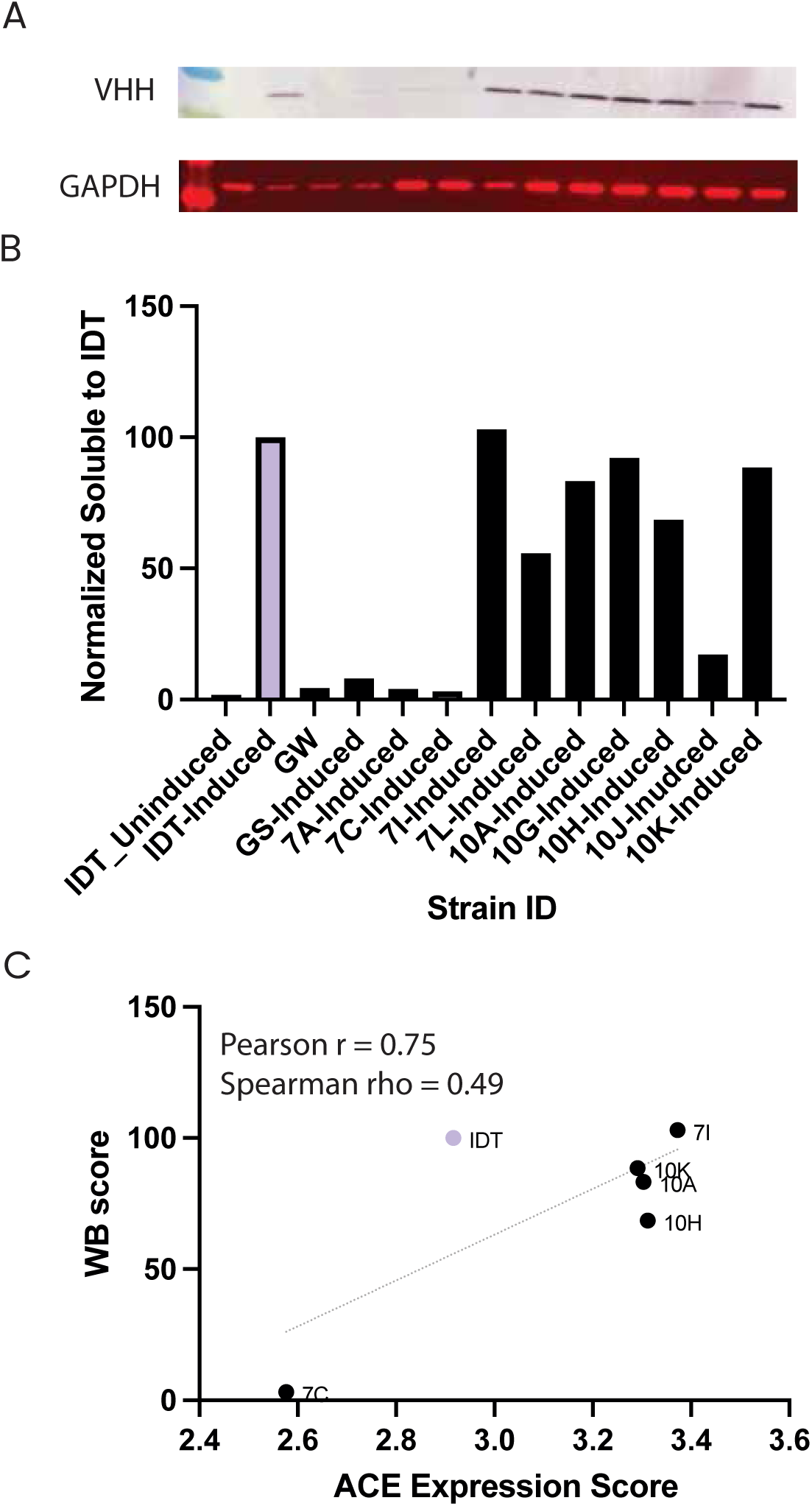
Validation of Anti-Her2 VHH expression scores via Western Blot. **(A)** Western blots of sampled strains from anti-HER2 VHH library. Blots show levels of VHH molecule and GAPDH solubility control. **(B)** Levels of soluble protein as measured by band density of Western Blots from A. Levels are calculated as density of VHH/GAPDH, then normalized to the IDT parent strain levels. **(C)** Correlation of strains from Western Blot in (A) with ACE derived expression scores for strains.

**Fig S14.**
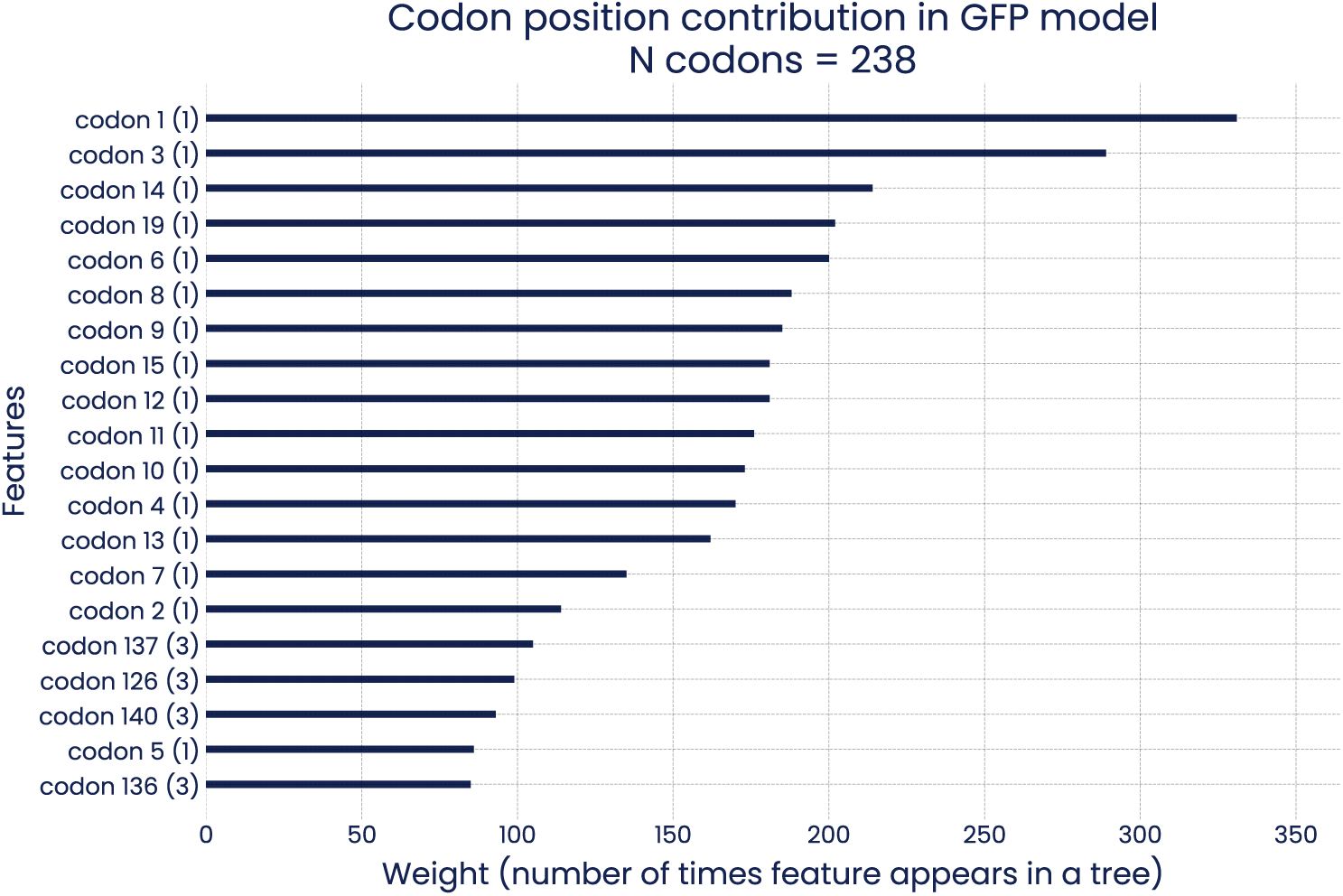
Codon importance in predicting GFP expression values in XGBoost model trained on one-hot encoded representations. The chart shows the top 20 most important features (codons) used by the XGBoost model to predict expression values. Each codon is numbered according to its position in the sequence and the number in brackets denotes its positional quartile, where 1 means the codon is found in the first fourth of the sequence and a 4 means the last fourth of the sequence.

**Fig S15.**
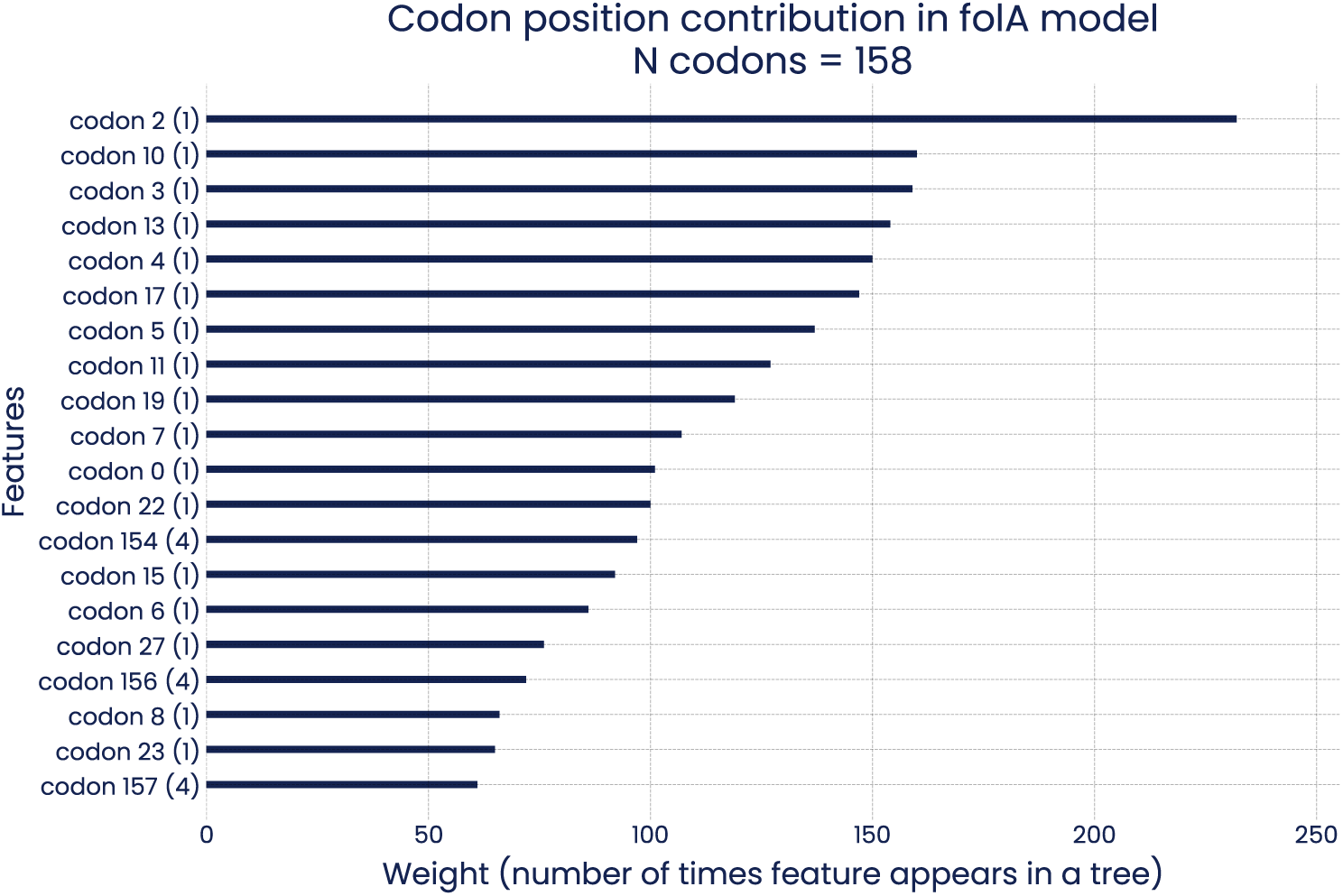
Codon importance in predicting *folA* expression values in XGBoost model trained on one-hot encoded representations. The chart shows the top 20 most important features (codons) used by the XGBoost model to predict expression values. Each codon is numbered according to its position in the sequence and the number in brackets denotes its positional quartile, where 1 means the codon is found in the first fourth of the sequence and a 4 means the last fourth of the sequence.

**Fig S16.**
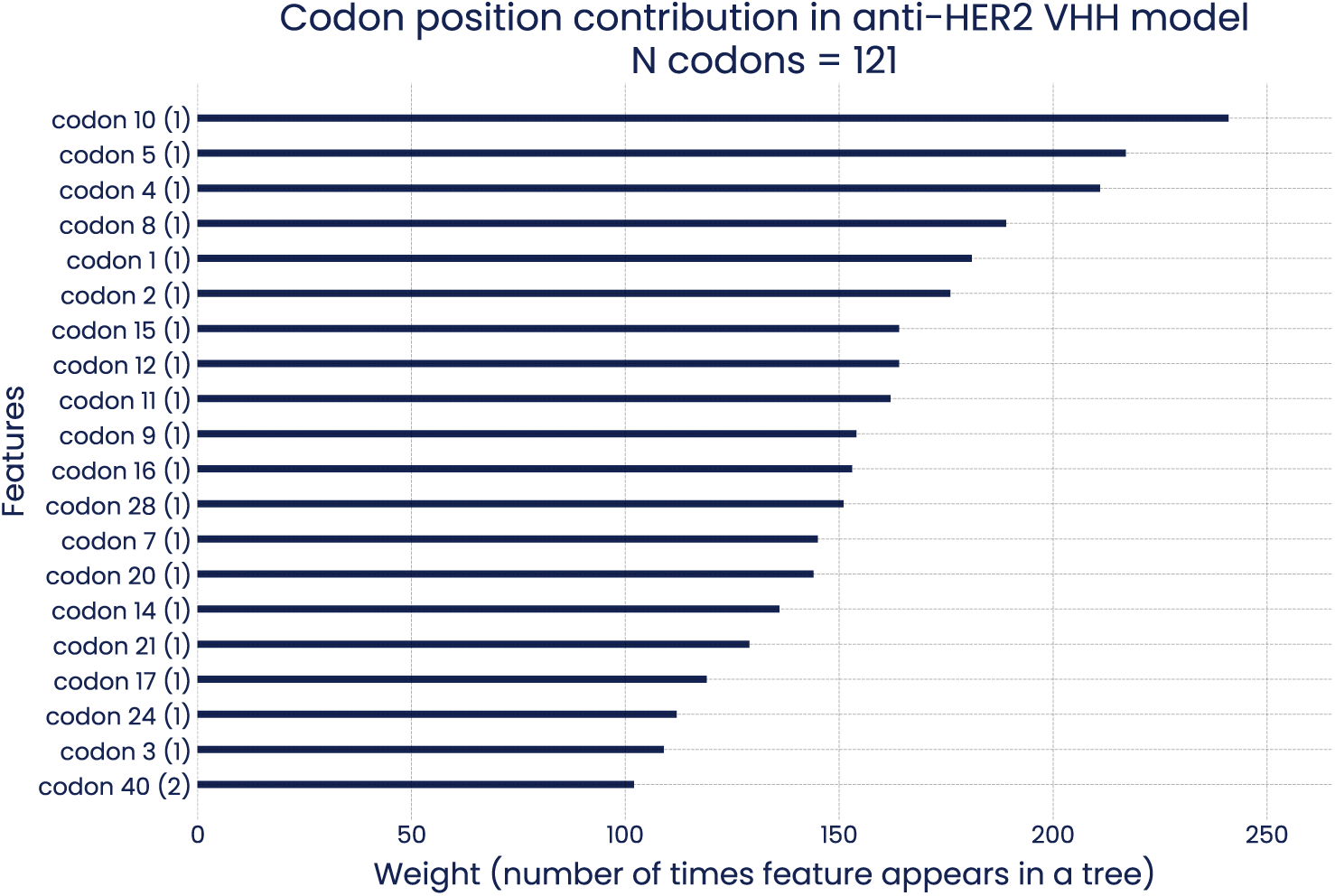
Codon importance in predicting anti-HER2 VHH expression values in XGBoost model trained on one-hot encoded representations. The chart shows the top 20 most important features (codons) used by the XGBoost model to predict expression values. Each codon is numbered according to its position in the sequence and the number in brackets denotes its positional quartile, where 1 means the codon is found in the first fourth of the sequence and a 4 means the last fourth of the sequence.

**Fig S17.**
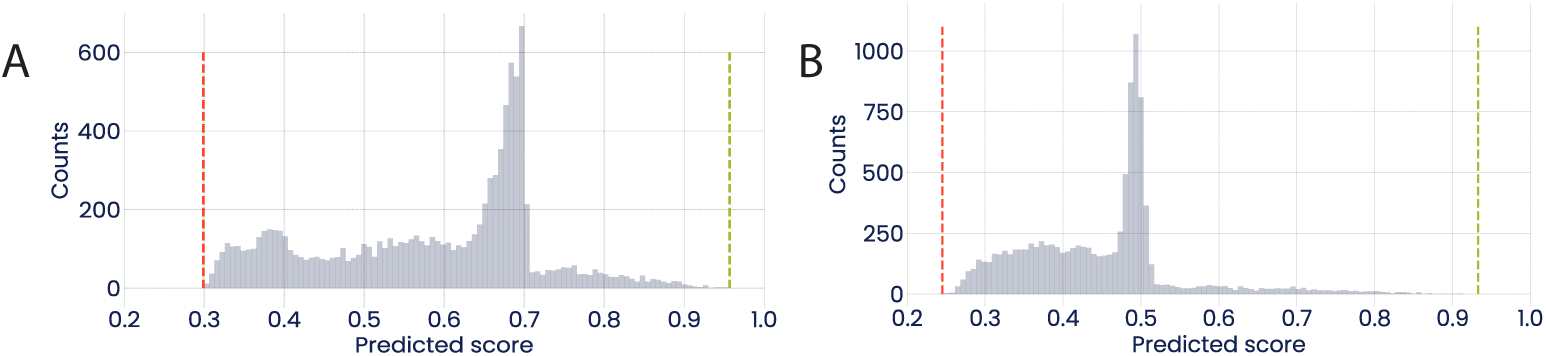
Distribution of randomly sampled and scored sequences for design of mCherry DNA sequences. **(A)** Distribution of a representative subset of sampled and scored sequences using the GFP model. Red dashed line represents the median GFP model score of the lowest 10 sampled variants. Green dashed line represents the median GFP model score of the highest 10 sampled variants.**(B)** Distribution of a representative subset of sampled and scored sequences using the ALL model. Red dashed line represents the median ALL model score of the lowest 10 sampled variants. Green dashed line represents the median ALL model score of the highest 10 sampled variants.

**Fig S18.**
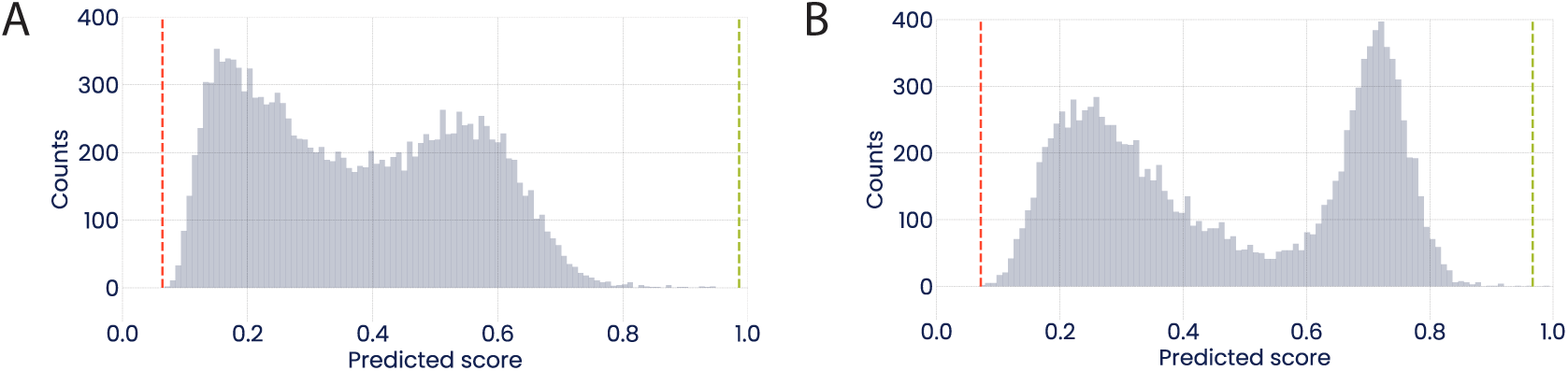
Distribution of randomly sampled and scored sequences for design of SARS-CoV-2 VHH DNA sequences. **(A)** Distribution of a representative subset of sampled and scored sequences using the VHH model. Red dashed line represents the median VHH model score of the lowest 10 sampled variants. Green dashed line represents the median VHH model score of the highest 10 sampled variants. **(B)** Distribution of a representative subset of sampled and scored sequences using the ALL model. Red dashed line represents the median ALL model score of the lowest 10 sampled variants. Green dashed line represents the median ALL model score of the highest 10 sampled variants.

**Fig S19.**
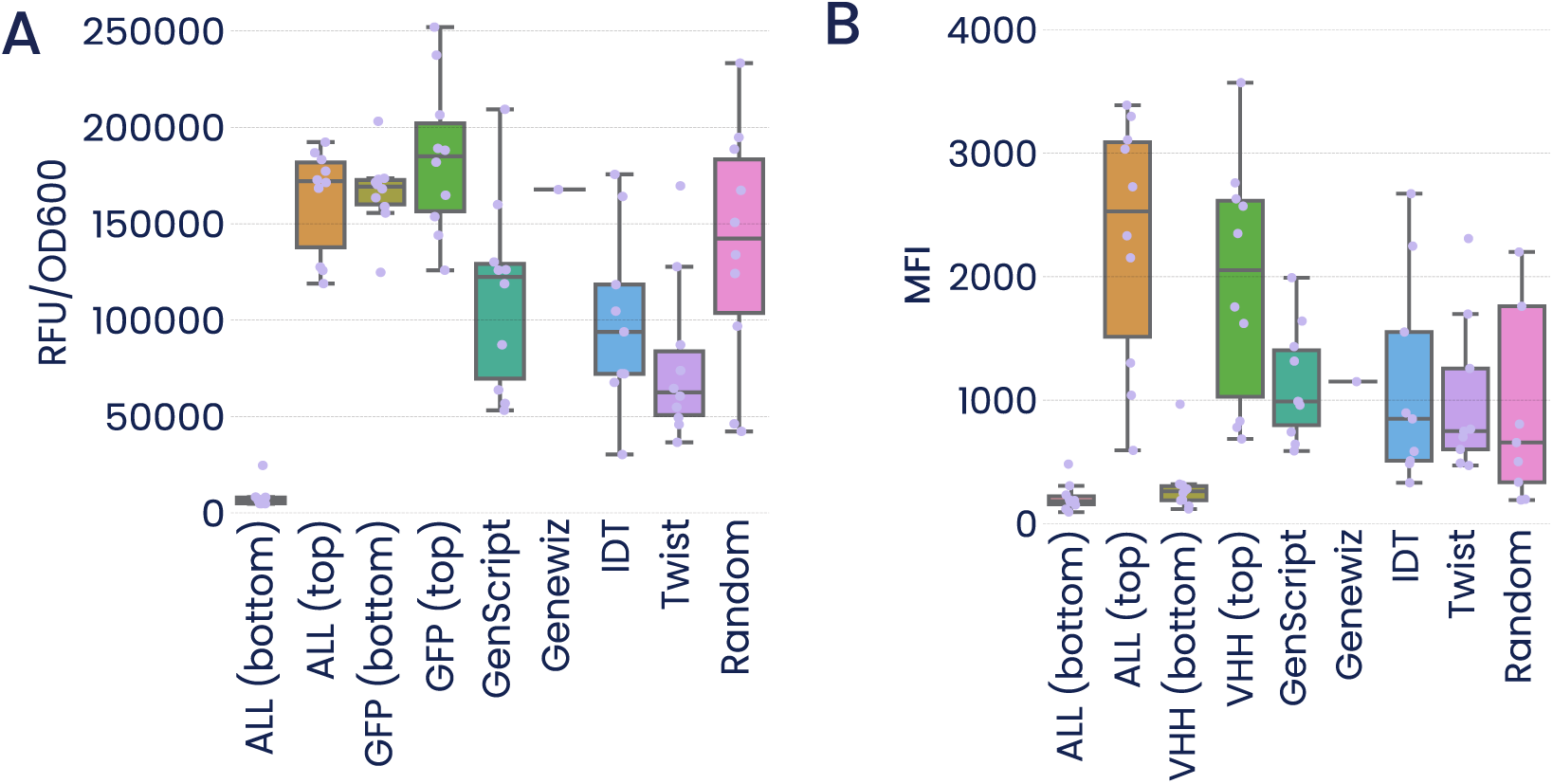
mCherry and anti-SARS-CoV-2 VHH model and commercial designs full data set. **(A)** Boxplot of all model and commercially designed sequences of mCherry. Note the high expression level of the GFP (bottom) 10 designs. **(B)** Boxplot of all model and commercially designed sequences of anti-SARS-CoV-2 VHH.

**Fig S20.**
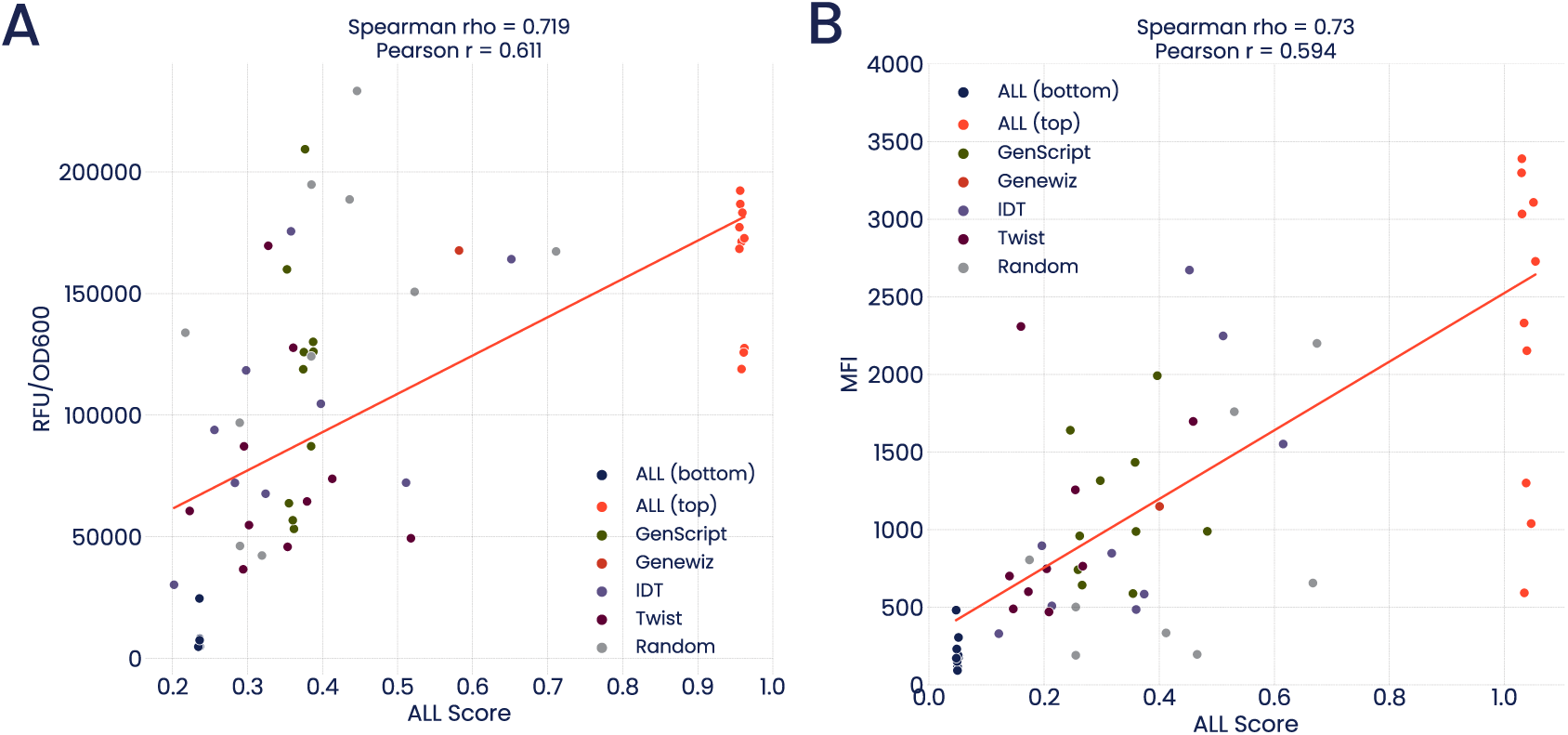
Correlation of ALL model scores from mCherry and anti-SARS-CoV-2 VHH variants tested in Figure 5. **(A)** Correlation plot of ALL model scores and functional expression fluorescence measurements of mCherry sequences from figure 5A. **(B)** Correlation plot of ALL model scores and ACE functional expression measurements of SARS-CoV-2 VHH sequences from figure 5C.

**Table S1.** Total possible codon sets for parental protein CDSs.

| Total possible CDSs for parental sequences. |  |  |
| --- | --- | --- |
| Protein | Total possible CDSs | Scientific Notation |
| **GFP** | 211571763348732056134479<br>734663327929687263573998<br>568424003830071633904161<br>158404976333163545734873<br>101171859390464 | 2.12e+110 |
| *folA* | 715452215168408987669254<br>286081230476835420164025<br>233822016678149236001970<br>5856 | 7.15e+75 |
| **Anti-HER2 VHH** | 150455113334121364441396<br>353542123355740550836658<br>190824046067712 | 1.50e+62 |
| **Anti-SARS-CoV-2 VHH** | 415978297346178748407572<br>638273262653951474953192<br>5659903225680101376 | 4.16e+66 |
| **mCherry** | 163249817398713005471849<br>779922894331418070510442<br>104016624597418269247531<br>506713785697315061925960<br>422535462912 | 1.63e+107 |

**Table S2.**
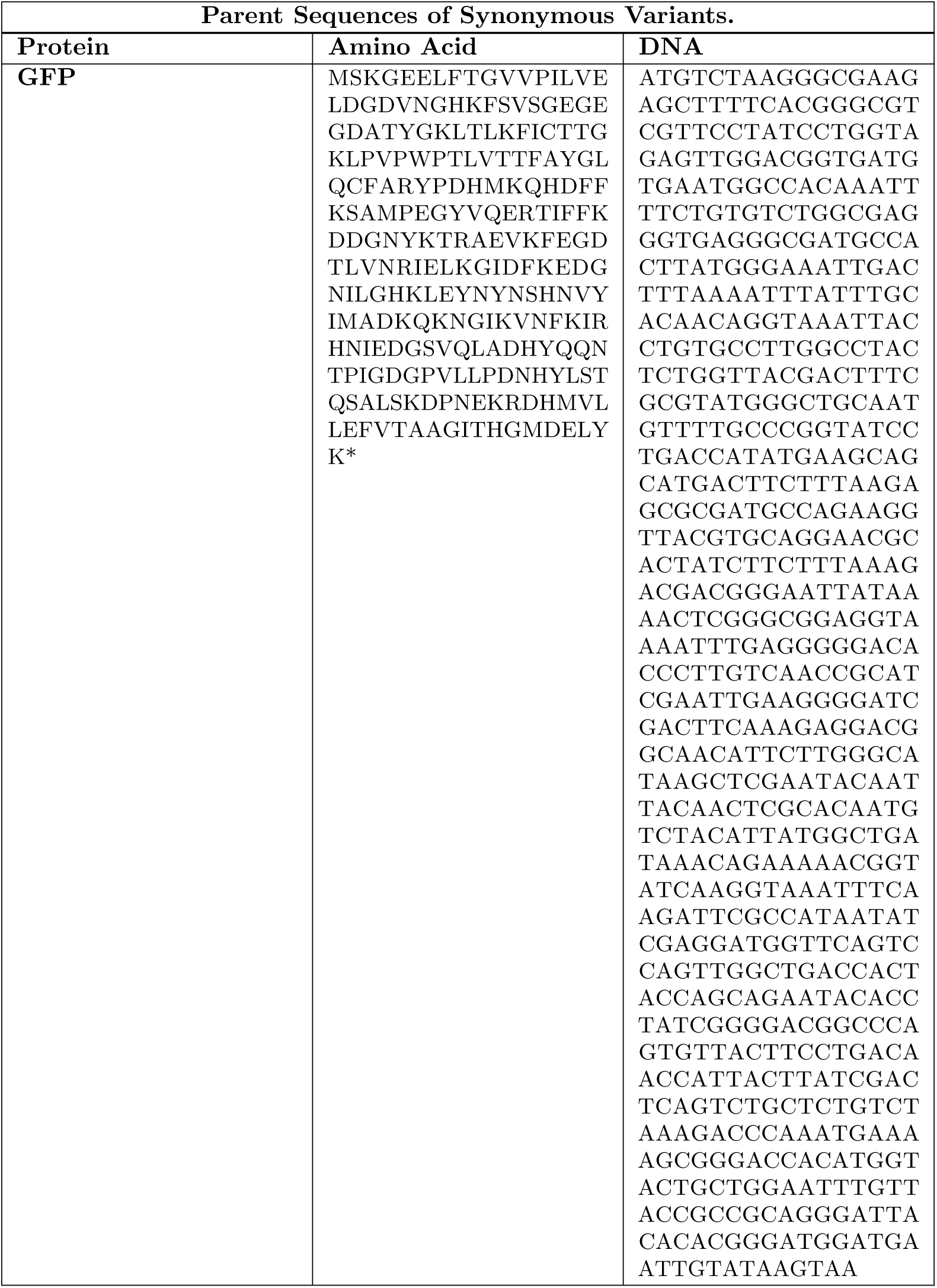
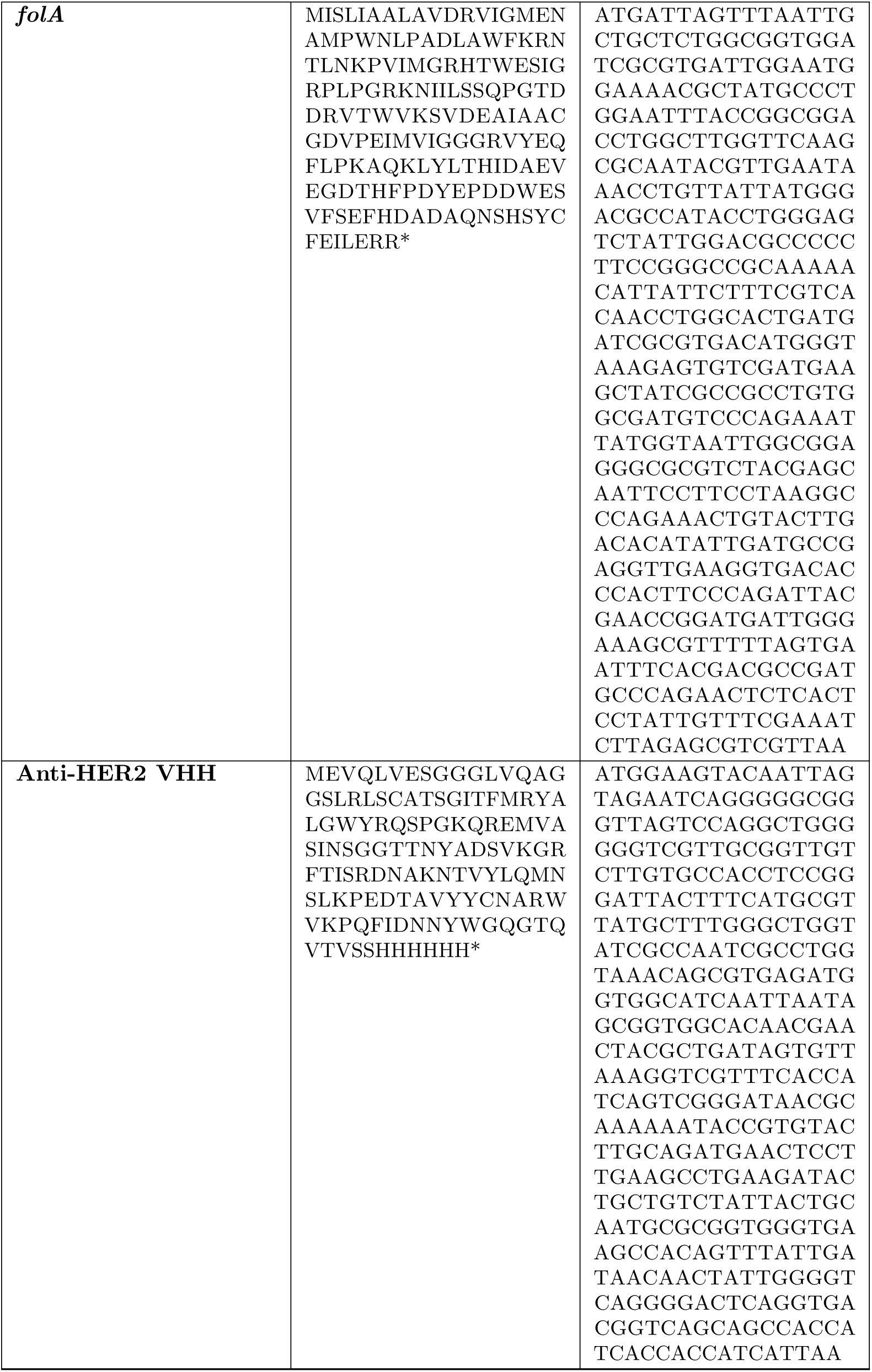
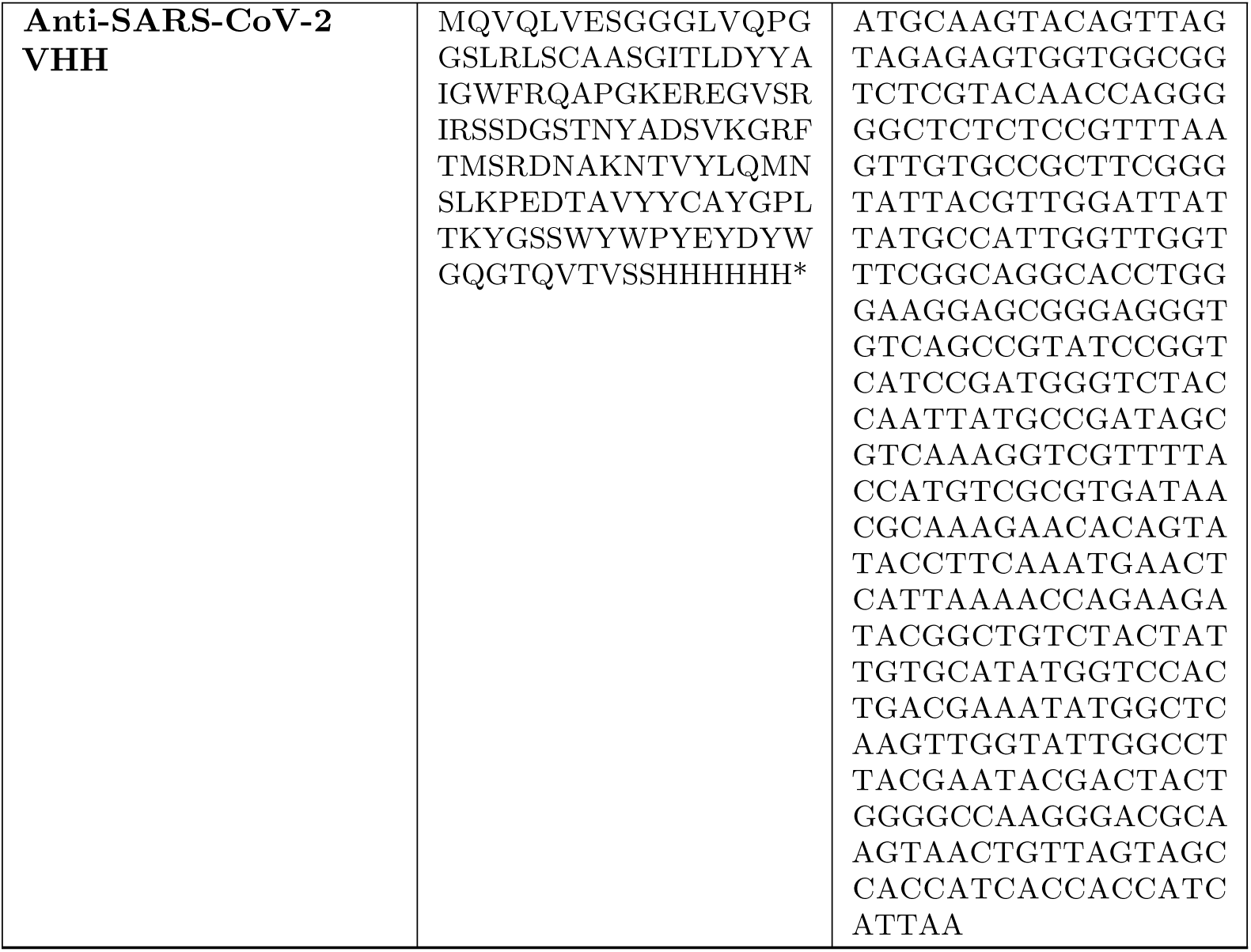
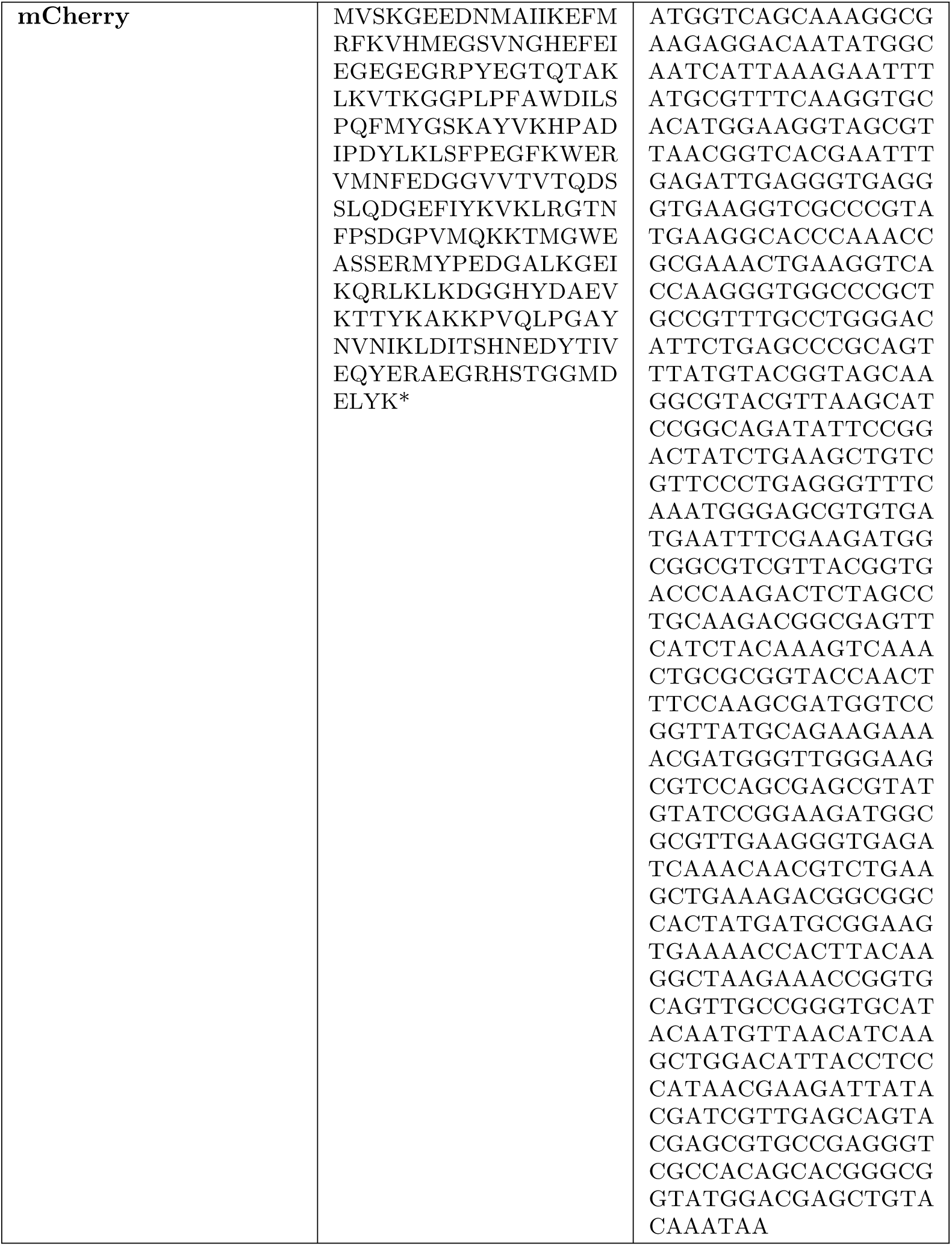
Parental sequences used to generate synonymous mutants.

**Table S3.** Number and percent of sequences that remained after each filtering step.

| Processing of Sequence Data for Expression Scores |  |  |  |  |  |  |
| --- | --- | --- | --- | --- | --- | --- |
| Dataset |  | Raw Sequences | Correct AA sequence |  | Over 10 reads |  |
| Target | Library | # | # | % | # | % |
| GFP | tile 1 | 1,318,099 | 862,168 | 65% | 42,480 | 3.22% |
| GFP | tile 2 | 1,587,238 | 1,074,099 | 68% | 45,152 | 2.84% |
| GFP | tile 3 | 452,072 | 264,379 | 58% | 28,778 | 6.37% |
| *folA* | Library A | 2,870,111 | 1,170,113 | 41% | 2,601 | 0.09% |
| *folA* | Library B | 3,547,929 | 1,541,237 | 43% | 3,145 | 0.09% |
| *folA* | Library C | 3,028,978 | 1,152,180 | 38% | 3,092 | 0.10% |
| *folA* | Library D | 4,324,141 | 1,903,028 | 44% | 3,811 | 0.09% |
| *folA* | Library E | 3,135,617 | 1,099,638 | 35% | 1,977 | 0.06% |
| *folA* | Library F | 3,709,504 | 810,249 | 22% | 1,497 | 0.04% |
| VHH | VHH | 3,698,925 | 2,097,177 | 57% | 16,279 | 0.44% |

**Table S4.** Spearman correlation (*ρ*) for fine-tuned and baseline models across holdout datasets (all p-values *<*0.01)

| Model | GFP data | folA data | VHH data |
| --- | --- | --- | --- |
| GFP full | 0.801 | 0.311 | 0.241 |
| GFP | 0.759 | 0.096 | 0.125 |
| *folA* | 0.347 | 0.899 | 0.428 |
| VHH | 0.137 | 0.024 | 0.782 |
| GFP+*folA* | 0.776 | 0.916 | 0.368 |
| GFP+VHH | 0.766 | -0.168 | 0.804 |
| *folA*+VHH | 0.629 | 0.905 | **0.837** |
| ALL | 0.788 | **0.918** | 0.818 |
| Pre-trained only | 0.269 | 0.014 | -0.433 |
| Finetuned only | 0.769 | 0.897 | 0.746 |
| XGBoost CO-T5 | 0.740 | 0.824 | 0.576 |
| XGBoost 1HE | **0.793** | 0.905 | 0.768 |

**Table S5.** Pearson correlation (r) for fine-tuned and baseline models across holdout datasets (all p-values <0.01)

| Model | GFP data | folA data | VHH data |
| --- | --- | --- | --- |
| GFP full | 0.854 | 0.217 | 0.181 |
| GFP | 0.808 | 0.054 | 0.057 |
| *folA* | 0.519 | 0.907 | 0.413 |
| VHH | 0.253 | -0.061 | 0.794 |
| GFP+*folA* | 0.830 | 0.910 | 0.340 |
| GFP+VHH | 0.824 | -0.127 | 0.815 |
| *folA*+VHH | 0.558 | 0.900 | **0.848** |
| ALL | **0.856** | **0.919** | 0.824 |
| Pre-trained only | 0.241 | 0.116 | -0.404 |
| Finetuned only | 0.835 | 0.900 | 0.745 |
| XGBoost CO-T5 | 0.760 | 0.788 | 0.555 |
| XGBoost 1HE | 0.843 | 0.898 | 0.768 |

**Table S6.** Performance of XGBoost baseline models trained with CO-T5 embeddings across the GFP (top), *folA* (middle), and anti-HER2 VHH (bottom) holdout datasets. A star (*) next to the Spearman and Pearson correlation coefficients denote statistical significance (p*<*0.01)

| Test dataset | Training dataset | Spearman $\rho$ | Pearson $r$ |
| --- | --- | --- | --- |
| GFP | GFP | 0.740* | 0.761* |
|  | *folA* | -0.144* | -0.174* |
|  | VHH | -0.179* | -0.133* |
| *folA* | *folA* | 0.824* | 0.788* |
|  | GFP | -0.261* | -0.221* |
|  | VHH | -0.046 | -0.004 |
| VHH | VHH | 0.576* | 0.555* |
|  | GFP | 0.167* | 0.155* |
|  | *folA* | -0.127* | -0.114* |

